# Matrix stress-relaxation and stiffness modulate oligodendrocyte differentiation

**DOI:** 10.1101/2025.10.09.680975

**Authors:** Eva D. Carvalho, Miguel R. G. Morais, Georgia Athanasopoulou, Marco Araújo, Sofia C. Guimarães, Stefano Pluchino, Cristina C. Barrias, Ana P. Pêgo

**Affiliations:** Instituto de Investigação e Inovação em Saúde (i3S), Universidade do Porto, Porto, Portugal; Instituto Nacional de Engenharia Biomédica (INEB), Universidade do Porto, Porto, Portugal; Instituto de Ciências Biomédicas Abel Salazar (ICBAS), Universidade do Porto, Porto, Portugal; University of Cambridge, Cambridge, United Kingdom

## Abstract

Impaired remyelination capacity, observed not only in demyelinating diseases of the central nervous system (CNS) but also in the aging brain, features a critical challenge in neural repair. Dysfunctional oligodendrocytes (OLs) are not able to efficiently restore myelin sheaths leading to increased axonal susceptibility to degeneration. Inhibitory cues to OL myelination have been described but strategies promoting remyelination keep failing.

Recently acknowledged as key regulators of cellular fate and behavior, physical properties are emerging as novel potential targets in remyelination. Yet, the full extent of this impact is unknown. So far, the viscoelastic properties of the brain have been overlooked, with studies assuming that it only possesses an elastic behavior.

Here, we engineered mechanically tunable 3D alginate hydrogels for culturing OLs. While being structurally biologically relevant, our matrices can be modelled in terms of elastic and viscoelastic properties. For the first time, we proved that, in addition to elasticity, viscoelasticity highly impacts the behavior of OLs. High stress-relaxation and shear moduli hydrogels lead to impaired differentiation, branching ability and metabolic activity of OLs, without visible effects on cellular viability. We showed that tuning alginate stress-relaxation properties while maintaining the stiffness triggers activation of mechanotransduction genes. Alterations were seen in genes involved in the focal adhesion kinase pathway as well as in the transcriptional factors Yap 1 and Taz and at the level of nuclear mechanotransduction (*Hdac1*).

Understanding how microenvironmental cues support OL viability and myelination might lead to the design of improved therapies promoting remyelination. Our tunable hydrogels are not only relevant to study OL mechanobiology but can also function as a relevant model to test novel therapeutic interventions in remyelination context.

## Introduction

Oligodendrocyte (OL) dysfunction and myelin loss are becoming increasingly recognized as defining features of many neurological diseases of the central nervous system (CNS). In addition, cognitive impairment observed in the aged brain is often attributed to the massive loss of the lipid-rich membrane layer that ensheaths axons[1]. In both scenarios, myelin clearance by CNS inflammatory cells is insufficient, hampering sheath repair. With time, remyelination led by OLs or oligodendrocyte precursor cells (OPCs) is also limited, and cannot assure the coverage of the full extent of myelin loss [2]. Damage to myelin results in increased axonal susceptibility to degeneration, as well as disturbances on signal propagation along neurons, causing serious cognition and motor problems culminating in life quality impairment. While the role of the cells and biomolecules (such as growth factors, surface ligands, and extracellular components) involved in these (de)myelination processes has been extensively studied, until now no single therapy has proved efficient in promoting remyelination [3].

OL function is highly modulated by their intricate and complex microenvironment. Physical properties as extracellular matrix (ECM) stiffness, topography, mechanical strain, and cellular confinement affect all stages of OL development (proliferation, migration, differentiation, and myelination) [4–12]. However, the majority of the models used in these studies primarily rely on two-dimensional (2D) biologically relevant surfaces and only take into account the substrate stiffness or the linear elastic modulus to characterize the mechanical environment [12, 13]. It is well known that cells interact with their surroundings, continually generating and maintaining traction forces with the environment. These forces, in turn, result in a time-dependent mechanical response known as viscoelasticity [14]. Recent findings have shown that the addition of a time constant will push mechanobiology studies to another level as elasticity and viscosity distinctly affect cellular behavior [14–16]. Although still scarce, research on the viscoelastic properties of the brain has shown that during development, aging or even in neurological diseases the viscoelastic properties of the brain dynamically change [17–20]. Nevertheless, the extent to which these properties affect OL demyelination and remyelination is poorly understood.

The process of OL differentiation and axonal myelin ensheathment involves extensive membrane and cytoskeleton remodeling, which in turn generates a series of forces between the OLs and the surrounding environment and/or the axons. However, until now there are no models that allow for the study of the viscoelasticity effect on OLs neither in 2D nor in 3D.

Here we propose the use of three-dimensional tunable alginate hydrogels to mimic the OPC/OL milieu and specifically study how mechanical properties affect the course of OL differentiation.

Being ultrasoft structures with a high-water content, hydrogels constitute one of the most biologic relevant matrices to mimic brain’s microenvironment. We have previously reported that alginate hydrogels are appropriate to culture neural cells, specifically astrocytes [21–23]. By introducing cell adhesive (RGD) and matrix metalloproteinases sensitive peptides (PVGLIG) we proved that astrocytes acquire a branched phenotype, and that the incorporation chemistry plays a role in cellular behavior [21]. Here we sought to use similar alginate hydrogels to culture OLs and, by tuning matrix elastic and viscoelastic properties, question the impact of the latter on OL differentiation processes.

For the first time, the combination of high stiffness and high stress-relaxation properties was described to have a negative impact on myelination. Additionally, both properties were found to regulate OL fate via distinct mechanisms. Stiffness caused visible changes in the Yap1/Taz transcriptional factors, while alterations in viscoelasticity strongly affected focal adhesion kinases related pathways and nuclear mechanotransduction, leading to transcriptional and epigenetic changes.

## Materials and Methods

### 1. Alginate functionalization with cell adhesive (RGD) and matrix metalloproteinases (MMP) sensitive peptides

All alginates used in the present study were derived from ultrapure high-molecular weight (HMW) sodium alginate (PRONOVA UP LVG, Novamatrix, FMC Biopolymers) with high guluronic acid content (68%), molecular weight (Mw) 244.1 ± 3.1 kDa and viscosity 20-200 mPa.s. The cell-adhesive RGD, GGGGRGDSP (Mw 758.74 g/mol) and the matrix metalloproteinase (MMP) sensitive GGYGPVG↓LIGK (Mw 1074.23 g/mol, the arrow indicates the cleavage site at the peptide bond between glycine, G, and leucine, L) peptides were purchased from GenScript.

#### 1.1 Grafting of RGD to alginate via carbodiimide chemistry

RGD was grafted to alginate via carbodiimide chemistry, as previously described [21, 24]. Alginate (1% wt/v, 200 mg) was dissolved in 2-(N-morpholino)ethanesulfonic acid (MES, Alfa Aesar) buffer (0.1 M MES, 0.3 M NaCl, dissolved in type II water and prepared protected from light), pH 6.5 for 1 h at room temperature (RT) (20 mL vial for 200 mg). The solution was transferred to an ice box and left under Argon environment for 1 min. N-hydroxysulfosuccinimide sodium salt (Sulfo-NHS, 0.054 mmol, Sigma) was poured down to the alginate solution and the system was immediately put under Argon atmosphere for 1 min. Ethyl-N′-(3-dimethylaminopropyl)carbodiimide hydrochloride (EDC.HCl, Sigma) (0.109 mmol) was dissolved in MES buffer and transferred to the alginate solution with the help of a syringe, in an inert atmosphere. After 10 min stirring, the peptide GGGGRGDSP (0.026 mmol) was dissolved in MES buffer and the solution was stirred overnight (o.n., 18 h) at RT in an inert atmosphere. The reaction was quenched with hydroxylamine hydrochloride (0.108 mmol, Sigma) and then transferred to a previously hydrated (20 min in type II H_2_O) dialysis membrane (MWCO 3500 membrane, Spectra/Por®, SpectrumLabs). Dialysis was performed for 2 days in decreasing concentrations of sodium chloride (128 nM – 21 nM NaCl, VWR, dissolved in 4 L type II H_2_O in polypropylene beakers) and for one day in type II H_2_O. Two water changes were performed in the first day (time interval 6 h) while three changes were performed in the following two days (time interval 4 h). Alginate grafted with RGD was then frozen at -80°C for at least 4 h and the lyophilized (for 2.5 days). Lyophilized alginate was weighted, and frozen at -20°C until further use. It is worth mentioning that a blank solution without the peptide was also performed following a similar protocol and used for quantification of peptide incorporation.

#### 1.2 Grafting of PVGLIG to alginate via carbodiimide chemistry or reductive amination

PVGLIG was either grafted to the alginate backbone through carbodiimide chemistry or reductive amination as previously described [21, 25, 26]. Carbodiimide coupling of PVGLIG was performed following a similar protocol as for RGD grafting. Briefly, 1% (wt/v) alginate (100 mg) was dissolved in 0.1 M MES buffer (0.1 M MES, 0.3 M NaCl, dissolved in type II water and prepared protected from light, pH 6.5) for 1 h, RT (20 mL vial for 200 mg). EDC.HCl (0.056 mmol) and sulfo-NHS (0.055 mmol) were added to the solution under Argon environment at 4°C. After 10 min stirring, the peptide GGYGPVG↓LIGK (0.015 mmol), previously dissolved in type II H_2_O and reacted with triethylamine (0.044 mmol, Riedel-de Haen) for 1 h at RT (without stirring), was added to the reaction under Argon conditions at 4°C. The reaction was stirred o.n. (18 h) at RT and afterwards quenched with hydroxylamine hydrochloride (0.055 mmol). Dialysis was carried out in the same manner as for RGD coupled via carbodiimide chemistry. A control reaction without the peptide was performed using the same methodology.

To engraft PVGLIG via reductive amination, alginate was previously oxidized as described elsewhere [21, 27]. Briefly, 1% (wt/v) alginate was dissolved in type II H_2_O for 1 h at RT (100 mL Schott for 600 mg) and then stirred for 10 min under ice. Sodium periodate (0.101 mmol for Alg Ox10 and 0.05 mmol for Alg Ox5, numbers following the “Ox” indicate the mol% of sodium periodate in the reaction, Sigma) was poured down to the alginate solution, protected from light. The tube that contained the sodium periodate was then washed with 100 μL type II H_2_O and added to the alginate solution. 2-propanolol (0.101 mmol for Alg Ox10 and 0.05 mmol for AlgOx5, VWR) was then added to the reaction to capture non-reacted radicals. The reaction was stirred for 10 min under ice and then o.n (18 h) at RT and protected from the light due to the oxidant light-sensitive properties. Afterwards, ethylene glycol (0.101 mmol for Alg Ox10 and 0.05 mmol for Alg Ox5, Merck) was added to the solution and stirred for 1 h at RT. The solution was then filtered through a 0.22 μm polyethersulfone (PES) filter (VWR). The product was purified by dialysis using a pre-hydrated (20 min, RT, type II H_2_O) MWCO 3500 membrane (Spectra/Por®, SpectrumLabs) membrane against type II H_2_O (4 L in polypropylene beakers) for 3 days, changing the water twice a day (8h interval). Solution was then frozen at -80°C and lyophilized.

After being oxidized, 10% (wt/v) alginate (100 mg) was dissolved in type II H_2_O and methanol (Sigma) at a ratio of 8.1:1 for 2h20 at RT (50 mL Schott for 100 mg alginate). GGYGPVG↓LIGK (0.015 mmol) was dissolved in type II H_2_O and reacted with triethylamine for 1h at RT without stirring. The peptide solution was added to the alginate under stirring and following the addition of the reducing agent. 2-Picoline-borane Complex (pic-BH_3_, Sigma-Aldrich) was dissolved in methanol (1.02 mmol) and added to the alginate solution. The final pH of the solution was adjusted to 5.2-5.5 using 1 M acetate buffer. Acetate buffer was prepared at pH 5.2 by firstly preparing 1 M sodium acetate (dissolved in type II H_2_O) and adjusting its pH to 5.2 with 1 M acetic acid (diluted in type II H_2_O). The 1 M acetate buffer was dropwise added to the alginate solution until the solution pH start dropping. A control solution without the peptide was performed to be used for peptide engrafting quantification purposes. After o.n. stirring (16 h) the solution was transferred to a hydrated dialysis membrane (MWCO 3500 membrane, Spectra/Por®, SpectrumLabs) and dialyzed against decrescent sodium chloride concentrations (128 mM to 0 mM, dissolved in 4 L type II H_2_O in polypropylene beakers) for 3 days (two changes with 6 h interval were performed in the first day, while three changes with 4 h time interval were conducted in the following two days). The alginate solution was frozen at -80°C in the morning of the next day (for minimum 4 h) and then lyophilized for 3 consecutive days. A control reaction without the peptide was performed to be used for peptide engrafting quantification purposes.

#### 1.3 Determination of the oxidation degree of alginate

To determine the oxidation degree of alginate we used a reported protocol with slight modifications [28]. Oxidized alginate (8 mg, performed in triplicate) was dissolved in 0.6 mL of 0.1 M acetate buffer (12 mL vials). The solution was stirred for 30 min at RT and afterwards tert-butyl carbazate (Sigma), at 0.315 M dissolved in 0.1 M acetate buffer, was added. After 1 h stirring at RT, sodium cyanoborohydride (NaBH_3_CN, Sigma) was weighted under a fume hood, dissolved in 0.1 M acetate buffer (0.297 M), and added to the alginate mixture. The solution was stirred o.n. (18 h) at RT and then transferred to a pre-hydrated dialysis membrane (MWCO 3500 membrane, Spectra/Por®, SpectrumLabs). Dialysis occurred for 3 days in decrescent sodium chloride concentrations (0.5-0 M, dissolved in 4 L type II H_2_O in polypropylene beakers) for 3 days. In the first day, two changes with 8 h time interval were performed (0.5 M NaCl), while in the remaining two days, three changes were made with a time interval of 6 h (day 2: 0.5 M (1x), 0.25 M (1x), 0.1 M (1x); day 3: 0 M (3x)). Solutions were then frozen at -80°C and then freeze-dried for 3 days. Samples were processed for nuclear magnetic resonance (^1^H NMR) evaluation by dissolution in deuterated water (13.3 mg/mL, D2O, Euriso-top) o.n. (18 h) at RT under agitation and then transferred to NMR tubes. ^1^H NMR spectra was recorded using a 400 MHz spectrometer AVANCE III (Bruker) using 32 scans. The spectra were processed using the software MestReNova (version 6.0). The oxidation degree was determined considering the peaks at δ= 4.51 ppm corresponding to the H1 proton from mannuronic residues of alginate and the one at δ=1.44 ppm belonging tert-butyl carbazate protons.

Supplementary Table S1. lists the oxidized alginate batches at 10% theoretical oxidation used in the context of the present study, with corresponding oxidation quantification and peptide incorporation.

#### 1.4 Quantification of peptide incorporation in alginate

The incorporation of RGD and PVGLIG in the alginate backbone was quantified using the colorimetric DC kit (BioRad). Briefly, 8 mg of corresponding alginate blanks control (alginate that submitted to a similar synthetic process but without peptides) was dissolved in 400 μL MiliQ H_2_O for 1 h at RT under stirring. In parallel, 1 mg (in triplicates) of alginate containing the peptides (Alg RGD or Alg PVG or Alg Ox PVG) was dissolved in 100 μL MiliQ H_2_O for 1 h at RT under stirring. To estimate the peptide concentration from the alginate solutions, firstly a calibration curve was performed. With a gel pipette, 300 μL of the blank solution were transferred to a new tube and diluted in 300 μL of MiliQ H_2_O (to achieve a 1% wt/v solution). From this solution, 80 μL were transferred to individual tubes (P1-P6). A 2 mg/mL peptide solution was also prepared by dissolution in 2 mg/mL in MiliQ H_2_O (without stirring or vortexing) which was then dissolved in the remaining 100 μL of the initially prepared blank solution (final peptide concentration 1 mg/mL, P7). Consecutive dilutions were performed from P7 to P1 by transferring 80 µL of solution, with P0 corresponding to the 1% (wt/v) blank solution without peptide.

With a gel pipette, 5 μL of the solutions contained in P0 to P7 tubes as well as from the alginate containing the peptide were transferred to a 96 transparent hydrophilic well-plate (Falcon, 353072), in triplicates. Following the manufacturer’s protocol, 25 μL of ralkaline copper tartrate solution (Reagent A) and 200 μL of Folin reagent (Reagent B) were added the wells. The mixture was incubated for 15 min at RT (protected from the light and without stirring) and then absorbance was read at λ = 750 nm using a micro-plate reader (Synergy MxTM, BioTek, USA).

### 2. Alginate hydrogels preparation

*In situ* crosslinked alginate hydrogels were prepared through internal gelation as previously described [21, 22, 29]. Briefly, stock solutions of 1.7% or 4% (wt/v) Alg HMW, Alg RGD or Alg PVG/Alg Ox PVG in 0.9% (wt/v) sodium chloride (pH 7.4) were prepared by o.n. stirring (2000 rpm) at RT. For cellular studies, only stock solutions of 1.7% (wt/v) alginate were filtered through a 0.22 μm polyvinylidene fluoride (PVDF) filter (Millex). Due to the high viscosity of the 4% (wt/v) alginate solutions it was impossible to filter these solutions.

For studies involving the optimization of an alginate hydrogel formulation for the growing of OLs, 1.7% (wt/v) solutions were mixed in adequate proportions to achieve alginate hydrogels containing 40 μM RGD (as previously optimized [21]) and different amounts of PVGLIG (200 and 400 μM). For studies involving the understanding of matrix mechanical properties impact on OL differentiation, RGD concentration was fixed to 40 μM and PVGLIG to 400 μM. For those studies the ratio of oxidized to non-oxidized was varied by filling alginate mixtures either with Alg HMW alginate (to achieve 2% Alg 30:70 and 3% Alg 20:80) or with oxidized alginate without PVGLIG (to achieve 2% Alg 60:40 and 3% Alg 60:40).

It is worth noting that 1% (wt/v) alginate hydrogels were obtained stock solutions at 1.7% (wt/v), whereas to achieve 2 and 3% (wt/v) alginate hydrogels, 4% (wt/v) alginate precursors were used.

Cell culture medium, containing or not cells, was added to the alginate solutions (for 1% (wt/v) alginate cell volume of 33% of the total final mixture volume; for 2% (wt/v) 43% and for 3% (wt/v) was 14%) and carefully homogenised using a 100 μL pipette gel (Gilson). An aqueous solution of calcium carbonate (CaCO_3_, Ca^2+/^COO^-^ molar ratio = 0.288, Fluka) was suspended in 0.9% (wt/v) autoclaved NaCl (initial concentration for 1% (wt/v) matrices = 130 mg/mL and for 2 and 3% (wt/v) = 285.4 mg/mL). It was then rapidly added to the alginate mixture and homogeneized. To increase the velocity of the crosslinking, glucone delta-lactone (GDL, Ca^2+^/GDL molar ratio = 0.125, Sigma) was rapidly dissolved in 0.9% NaCl (initial concentration for 1% (wt/v) matrices = 205.7 mg/mL and for 2 and 3% (wt/v) = 451.7 mg/mL). After filtering through a 0.22 μm PES filter (VWR), GDL was added to the alginate solution. GDL is responsible for the acidification of the alginate mixture and increase solubility of Ca^2+^ allowing a faster crosslinking of the polymer chains. Afterwards, the alginate mixture was pipetted (20 μL) onto wells of a non-treated 48 well plate (Falcon) and crosslinking occurred at 37°C, 5% CO_2_ for 30 min. Final CaCO_3_ concentration in the alginate mixtures for 1% (wt/v) matrices was equal to 1.02 mg/mL and for 2 and % (wt/v) equal to 2.04 mg/mL. Final GDL concentration in the alginate mixtures for 1% (wt/v) matrices was equal to 14.52 mg/mL and for 2 and % (wt/v) equal to 29.04 mg/mL.

### 3. Micropillars preparation

Polydimethylsiloxane (PDMS) micropillars were produced by casting the polymer into silicon molds, produced by eletron-beam lithography and deep reactive ion etching. PDMS structures were obtained by mixing Silastic® T-2 Translucent Base (viscosity 50,000 mPa.s) with the Curing Agent (Dow Corning) in a 10:1 base-to-curing agent ratio. Micropillars were cured at 60°C for 1 h 15 min and afterwards peeled off from the mold, cut out, immersed in 2-propanol and washed with an increasing gradient of ethanol:2-propanol (0:100, 50:50, 100:0) for a complete substitution of 2-propanol for ethanol. Then, micropillars were critical point dried (Polaron Range CPD7501, Quorum Technologies) using 7 cycles of flush (at 4°C, 800 psi) to substitute ethanol for liquid CO_2_. Micropillars of 5 μm diameter, interspaced 30 μm and with a height of 10 μm were used in this study.

### 4. Rheological properties of alginate hydrogels

#### 4.1 Shear modulus determination

Shear moduli of alginate hydrogels were determined by rheology using a Kinexus pro Rheometer (Malvern Instruments Ltd.). Alginate hydrogels with a diameter of 4 mm (cut with a puncher) were loaded on the bottom geometry plate (4 mm parallel-plate geometry) and oscillation tests were performed at 37°C in a humidified environment, with 30% compression (in relation to hydrogels’ height, previously estimated with a caliper). Measurements were done 24 h after hydrogel polymerization and the pH of the medium was equilibrated by immersing gels in Dulbecco’s Modified Eagle Medium (DMEM) Glutamax^TM^ High Glucose containing 25 mM N-2-hydroxyethylpiperazine-N-2-ethane sulfonic acid (HEPES) (Biowest), pH 7.5 for at least 30 min. The linear viscoelastic region (LVR) was determined using amplitude sweep tests at a constant oscillation frequency of 0.1 Hz and strains in varying from 0.1-100% (10 points per decade). Once the LVR was defined, oscillation frequency sweep tests were conducted. For frequency sweep tests, hydrogels were submitted to oscillations in the range of 0.01-5 Hz, at a constant strain of 1.5% (10 points per decade). Values of complex shear modulus (G*) elastic shear moduli (G’), viscous shear moduli (G’’) and phase angle (δ) were determined from the frequency tests, with a tolerance range of around the median value of ± 10%.

Mesh sizes (ξ) of the hydrogels were indirectly estimated from shear modulus measurements as previously shown by others [30]. The average mesh size is the distance between two crosslinking points and can be predicted from the equation:

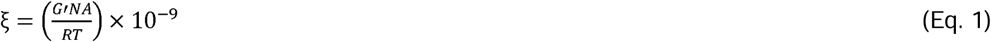

where NA is the Avogrado number (6.022 x 10^23^), R is the gas constant (8.314 J/K mol) and T is the absolute temperature (37°C = 310 K).

#### 4.2 Stress-relaxation measurements

The stress-relaxation properties of the alginate hydrogels were also measured using a Kinexus pro Rheometer (Malvern Instruments Ltd.). After 24 h of culturing, alginate hydrogels were immersed in DMEM with 25 mM HEPES. Hydrogels were compressed to 10% strain with a deformation rate of 3 min. The strain was held constant while the load was recorded as a function of time and 50 samples per decade were considered.

### 5. Cell cultures

#### 5.1 Mixed glial cell (MGC) cultures

All procedures involving animals and their care were performed in agreement with institutional ethical guidelines (IBMC/INEB/i3S), the EU directive (2010/63/EU), and the Portuguese law (DL 113/2013). Experiments described here had the approval of Portuguese Veterinary Authorities. Animals had free access to food and water, being kept under a 12-h light/12-h dark cycle.

Primary cultures of OPCs were obtained as previously described [22, 31, 32]. Briefly, P0-P2 Wistar Han rats were sacrificed by decapitation and brain removed and dissected in Hank’s Balanced Salt Solution without calcium and magnesium (HBSS, Sigma) supplemented with 2% (v/v) penicillin/streptavidin (P/S) on ice. Isolated cortices were digested in HBSS without calcium or magnesium supplemented with trypsin (0.0025% (w/v)) and 0.001 mg/mL DNAse I (Applichem LifeSciences) for 15 minutes at 37°C. Dissociated cortices were cultured in 10 µg/mL poly-D-lysine (PDL, Mw >300,000 Da, Merck A003E) coated 75 cm^2^ flasks (30 min, 37°C, and 5% CO_2_) and maintained in DMEM supplemented with 10% (v/v) heat-inactivated (30 min, 56°C) foetal bovine serum (FBS, Sigma) and 1% (v/v) P/S (37°C and 5% CO_2_). When confluence was reached (∼12 days) the flasks were submitted to a pre-shake (200 rpm, 2 h, 37°C) to detach microglia. Following a 2 h incubation period (37°C, 5% CO_2_), MGCs were submitted to shake-off (200 rpm, o.n, 37°C), resulting in detachment of OPCs and remaining microglia. Medium containing detached OPCs and microglia was transferred to non-coated and non-treated Petri dishes (90 mm in diameter) and incubated for 2 h (37°C, 5% CO_2_). Suspension medium with OPCs was passed through a 40 μm nylon cell strainer (Falcon) to remove astrocyte clusters and centrifuged for 10 min, 500 x g, at RT. Cells were then resuspended, and live cells counted using the Trypan blue assay.

#### 5.2 Oligodendrocytes cultures in alginate hydrogels and micropillars

OPCs were embedded within alginate hydrogels at a final concentration 2.25 x 10^6^ cells/mL (45 000 viable cells (trypan blue assay) in a 20 µL hydrogel) and culture was maintained until maximum of 14 days. OPCs were maintained in OL SATO medium which was composed by DMEM medium supplemented with apo-transferrin (0.1 mg/mL, Merck T2036), bovine serum albumin (BSA) (0.1 mg/mL, NZYTech MB04601), putrescin (16 µg/mL, Sigma P7505), progesterone (60 ng/mL, Sigma P8783-), thyroxine (40 ng/mL, Merck T1775), sodium selenite (40 ng/mL, Sigma 214485-), triiodo-L-thyroxine (30 ng/mL, Merck T1775), human insulin (5 µg/mL, Sigma I9278), 0.5% heat-inactivated (30 min at 56°C) FBS, and 1% (v/v) P/S.

For studies involving alginate hydrogels embedding a micropillar array, PDMS structures were plasma treated using the Diener Electronic Zepto Plasma Cleaner, at a low pressure (0.8 mbar), 30% intensity for 3 min. Immediately after, PDMS was coated with a mixture of poly-L-lysine (PLL, Mw 30,000-70,000, pH 6.5, final concentration 50 μg/mL Sigma-Aldrich P2636) and bovine serum albumin (diluted in type II H_2_O, pH 7.4, initial concentration 1 mg/mL) in a ratio of 5:4:1 of PLL: type II H_2_O : BSA. It is worth noting that the preparation of this solution required first the addition of PLL, then type II H2O and finally BSA carefully to avoid precipitates formation. The addition of BSA (a negatively charged protein) to PLL was seen to be needed to avoid hydrogel separation from the micropillars after crosslinking. Coating was performed o.n. at 37°C, 5% CO_2_. After washing twice with sterilized type II water, PDMS micropillars were coated with laminin 211 (Biolamina LN211, 10 μg/mL) and incubated for 2 h (37°C, 5% CO_2_). No washing was performed after laminin coating. Alginate hydrogels containing OPCs (final concentration 4.50 x 10^6^ cells/mL, 90 000 cells in a 20 µL hydrogel) were pipetted onto the micropillars and crosslinking of the alginate occurred for 30 min (37°C, 5% CO_2_). Cultures were performed in tissue-culture 12-well plates (Falcon) (total medium volume of 1.7 mL). Cell culture purity was determined above 70% as previously described [22].

### 6. Biocompatibility of alginate hydrogels

#### 6.1 Cellular metabolic activity (resazurin assay)

Metabolic activity was inferred by the measurement of resorufin fluorescence. Briefly, hydrogels containing OPCs were incubated with resazurin solution (10% (v/v), in OL SATO medium) for 3 h (37°C, 5% CO_2_). Supernatant medium was transferred to black 96-well plates and fluorescence measured at 530/590 nm in a multi-plate reader Synergy Mx (BioTeK® Instruments, GenX software).

#### 6.2 Cell viability (live-dead assay)

At a defined time point cells were washed with pre-warmed (37°C) tris-buffered saline-calcium chloride 15 mM solution (TBS-CaCl_2_ pH 7.4) for 5 min (37°C, 5% CO_2_) and incubated with a cell-permeant dye calcein-AM (0.25 μg/mL, λ_emission_ 517 nm/λ_excitation_ 484 nm, Promega C1430) for 2 h (37°C, 5% CO_2_). Cells were washed with TBS-CaCl_2_ 15 mM followed by a 5 min incubation at RT with propidium iodide solution (PI, 0.25 μg/mL, λ_emission_ 617 nm/λ_excitation_ 535 nm, Sigma-Aldrich), and Hoechst 33342 (10 μg/mL, λ_emission_ 497 nm/λ_excitation_ 361 nm, Molecular Probes). The excess of PI and Hoechst was removed by rinsing the hydrogels with TBS-CaCl_2_ 15 mM and immediately imaged under a confocal fluorescence microscope (37°C, 5% CO_2_).

### 7. Immunocytochemistry (ICC)

Cells were washed with pre-warmed 15 mM TBS-CaCl_2_ pH 7.4 and fixed with 4% (wt/v) microtubule-protecting paraformaldehyde solution (MP-PFA) for 20 min at RT. MP-PFA consists of 4% (wt/v) PFA (Merck Milipore), 65 mM PIPES (Milipore), 25 mM HEPES (Sigma-Aldrich), 10 mM EGTA (Sigma-Aldrich), 3 mM MgCl_2_ (Sigma-Aldrich) diluted in 15 mM TBS-CaCl_2_ pH 7.4 [33]. OLs were further permeabilized for 75 min in 2.5% (v/v) Triton X-100 solution diluted in 15 mM TBS-CaCl_2_ pH 7.4 (Sigma) under gentle agitation at RT. Afterwards, cells were blocked with 15 mM TBS-CaCl_2_ pH 7.4 containing 3% (v/v) BSA for 1 h at RT under gentle agitation. Primary antibodies were diluted in 15 mM TBS-CaCl_2_ pH 7.4 containing 1% (v/v) BSA and 0.5% (v/v) Triton X-100 and incubated o.n. in a humid chamber at 4°C, under gentle agitation. The following primary antibodies were used: monoclonal rat anti-myelin basic protein (anti-MBP) (1:100, Bio-Rad Laboratories, MCA409S) and monoclonal mouse anti-α tubulin (1:800, Sigma-Aldrich, T5168). After o.n incubation, hydrogels were washed three times with a solution of 0.05% Triton X-100 diluted in 15 mM TBS-CaCl_2_ pH 7.4 for 30 min, under gentle agitation. Secondary antibodies Alexa Fluor™ 647 anti-rabbit and 488 anti-rat (1:500, ThermoFisher Scientific, A-11008 and A21208, respectively) diluted in 15 mM TBS-CaCl_2_ pH 7.4 containing 3% (v/v) BSA and 0.5% (v/v) Triton X-100 were applied for 5 h at RT. Subsequently cells were washed twice with a solution of 0.05% Triton X-100 diluted in 15 mM TBS-CaCl_2_ pH 7.4 for 30 min, under gentle agitation and afterwards treated for nuclear counterstaining at RT for 15 min with Hoechst 33342 (1:500, λ_emission_ 497 nm/λ_excitation_ 361 nm, Molecular Probes, H3570). Hydrogels were washed again twice with 0.05% Triton X-100 diluted in 15 mM TBS-CaCl_2_ pH 7.4 for 30 min, under gentle agitation and stored at 4°C in 15 mM TBS-CaCl_2_ pH 7.4 containing 0.01% sodium azide (Sigma) until further imaging.

For Yap1 studies, a similar protocol was followed with slight modifications in the duration of the incubations and concentrations of some agents. Permeabilization and blocking were performed together during 24 h at 4°C in a solution of 0.3% (v/v) Triton X-100 and 3% (v/v) BSA diluted in 15 mM TBS-CaCl_2_ pH 7.4 followed by primary antibody incubation o.n. at 4°C, under gentle agitation. Primary antibodies used were mouse anti-Yap1 (1:100, Santa Cruz Biotechnology, SC-101199) and monoclonal rat anti-MBP (1:100, Bio-Rad Laboratories, MCA409S) and were incubated in a solution containing 1% (v/v) BSA and 0.15% (v/v) Triton X-100. Secondary antibodies (1:500) were applied for 2 days at 4°C and were the following: Alexa Fluor™ 647 anti-rabbit and 594 anti-rat (A21209). Alexa Fluor™ 488 Phalloidin (λ_emission_518 nm/λ_excitation_495 nm, ThermoFisherScientific, A12379) was diluted in 15 mM TBS-CaCl_2_ pH 7.4 (1:200) and hydrogels were incubated for 5 h at RT, under gentle agitation. Nuclear staining was performed as explained before.

For fluorescence visualization purposes, hydrogels were transferred, with the help of a spoon, to 35-mm ibidi dishes (ibidi 80136), turned upside down and a humidified environment (TBS-CaCl_2_ drops around the ibidi) was created to avoid hydrogel drying.

### 8. Confocal laser scanning microscopy

For the live-dead assays, OLs in alginate hydrogels were imaged using a TCS SP5 laser confocal inverted microscope (Leica Microsystems). Images were acquired using a HCX PL APO CS 10.0x0.40 DRY UV with a resolution of 16 bits and in a sequential mode. A frequency of 700 Hz was applied with a unidirectional scan, a frame average of 3 and a line average of 2. An area of 911.76 x 911.76 μm (1024 x 1024, zoom factor 1.7) and a z-step size of 5 μm (z-size of 115 μm) were applied. Calcein was excited with a 488 nm laser line with a laser power of 10% (set at 20% Argon laser power), a gain of 513 and an offset of 0. Emission light was collected on a Leica PMT detector with a collection window of 497-556 nm. PI was excited with a 594 nm laser line with a laser power of 9%, a gain of 713 and an offset of 0. Emission light was collected on a Leica PMT detector with a collection window of 604 – 642 nm. Hoechst was excited with a 405 nm laser line with a laser power of 23%, a gain of 718 and an offset of 0. Emission light was collected on a Leica PMT detector with a collection window of 427-494 nm. The pinhole size was 56.6 μm, calculated at 1 airy unit (AU) for 580 nm emission.

For image of fixed samples, a TCS SP8 confocal laser scanning microscope (Leica Microsystems) was used. For evaluation of MBP positive cells and OL morphology, images were acquired using a HC PL APO CS2 40x/1.10 WATER, with a resolution of 16 bits and in a sequential mode. A resonant frequency of 8000 Hz was applied with a bidirectional scan, a frame average of 1 and a line average of 6. An area of 232.5 x 232.5 μm (512 x 512 zoom factor 1.25) and a z-step size of 1.5 μm (z-size of 245 μm) were applied. 30 regions per hydrogel were acquired. MBP labelled with Alexa Fluor 488 nm was excited with an Argon laser with a laser power of 1.82% (set at 10% Argon laser power), a gain of 10 and an offset of 0. Emission light was collected on a Leica HyD 3 detector with a collection window of 498 – 572 nm. α tubulin labelled with Alexa Fluor 647 nm was excited with a HeNe 633 nm laser line with a laser power of 1.69% a gain of 10 and an offset of 0. Emission light was collected on a Leica HyD4 detector with a collection window of 643 – 726 nm. Hoechst was excited with a 405 nm laser line with a laser power of 10%, a gain of 752 and an offset of 0. Emission light was collected on a Leica PMT detector with a collection window of 416 – 479 nm.

For evaluation of Yap1 location, images were acquired using a HC PL APO CS2 40x/1.10 WATER, with a resolution of 16 bits and in a sequential mode. A frequency of 700 Hz was applied with a bidirectional scan, a frame average of 1 and a line average of 6. An area of 227.94 x 227.94 μm with a pixel size of 440 nm (512 x 512 zoom factor 1.28) and a z-step size of 2.5 μm (z-size of 40 μm) were applied. 9 regions per hydrogel were acquired. MBP labelled with Alexa Fluor 488 nm was excited with an Argon laser with a laser power of 4% (set at 10% Argon laser power), a gain of 22.8 and an offset of -0.01. Emission light was collected on a Leica HyD detector with a collection window of 498 – 572 nm. Yap1 labelled with Alexa Fluor 647 nm was excited with a HeNe 633 nm laser line with a laser power of 1.7% a gain of 31 and an offset of 0. Emission light was collected on a Leica HyD4 detector with a collection window of 643 – 726 nm. F-actin labelled with Alexa Fluor 488 Phalloidin was excited with an Argon laser with a laser power of 20% (set at 10% Argon laser power), a gain of 834.4 and an offset of -0.29. Emission light was collected on a Leica PMT detector with a collection window of 498 – 572 nm. Hoechst was excited with a 405 nm laser line with a laser power of 3%, a gain of 623.7 and an offset of -0.21. Emission light was collected on a Leica PMT detector with a collection window of 416 – 479 nm. The pinhole size was 77.2 μm, calculated at 1 airy unit (AU) for 580 nm emission.

### 9. Image segmentation and analysis

To assess MBP cell number, volume of MBP and sphericity, images were firstly converted from a. lif to a .tiff format using the ImageJ (version 1.52u 17 March 2020, https://imagej.nih.gov/ij/notes.html) [34] macro “LIFtoTIFF.ijm” (**Supplementary Pipeline 1**). Briefly, channels from the image were split and a 3D Gaussian blur filter of sigma 1 for channel 1 (correspondent to Hoechst labelling) and sigma 2 for channel 2 (correspondent to MBP labelling) were applied. Images were then saved in a .tiff format. Before proceeding to image segmentation and feature extraction, all images previously saved were included in a single folder using the Python 3 (https://www.python.org) script “fileinfolder.py” (**Supplementary Pipeline 2**). Subsequently, images were transferred to IMARIS (version 9.6.1, https://imaris.oxinst.com) and converted in .ims format. Nuclei were identified from Hoechst images by using the “Spots” tool to detect objects with an estimated diameter of 6 μm and a “quality” filter parameter above 2432. To segment OLs, the “Cells” tool was used. “Cells” were detected using a cell smooth filter width of 0.227 µm and a background subtraction width of 0.909 µm. The threshold was manually set to 264.128 and cells were filtered based on the number of voxels (above 2432). Positive cells for the MBP marker were estimated based on the detection of “Vesicles type A” from the segmented “Cells” containing the same parameters as the ones set for nuclei detection. Excel files containing statistics for “Spots” and “Cells” were retrieved from the software and then organized using the “restructIMARIS” pack created using Python 3 (https://www.python.org) (**Supplementary Pipeline 3**).

### 10. RNA extraction, cDNA synthesis and quantitative real-time polymerase chain reaction (qRT-PCR)

OPCs/OLs were retrieved from alginate hydrogels by firstly washing hydrogels with pre-warmed 15 mM TBS-CaCl_2_ pH 7.4 for 5 min (37°C, 5% CO_2_). At least 8 hydrogels from the same condition were pooled together and gels incubated with trypsin-ethylenediaminetetraacetic acid (EDTA) 50 mM for 7 min at 37°C, 5% CO_2_. For 1% (wt/v) alginate hydrogels 150 µL of trypsin-EDTA was added, while for 2% (wt/v) alginate hydrogels 300 µL and for 3% (wt/v) 450 µL of the solution was added. The action of the trypsin was stopped with at the least the same volume of OL SATO, solutions were homogenized and centrifuged for 5 min at 10 000 rpm (RT). Supernatants were discarded and the pellet washed with pre-warmed (37°C) phosphate buffer saline (PBS) pH 7.4. After centrifugation for 5 min at 10 000 rpm (RT), cells were lysed using lysis buffer (300 µL) from the Quick-RNA MiniPrep kit (Zymo Research R1050). Total RNA was then extracted according to the manufacturer’s recommendations. RNA concentration was estimated using the NanoDrop 1000 Spectrophotometer (ThermoFisher Scientific). Afterwards, cDNA was synthesized from 100 ng of RNA using the NZY First-Strand cDNA kit (NZY, MB12501) following the manufacturer’s protocols. Quantitative real-time PCR was performed on CFX 384 (Bio-Rad) in triplicates using the iTaq Universal SYBR Green Supermix (Bio-Rad) composed by 0.25 μL for each primer (final concentration 25 nM), 5 μL of iTaq, 1 μL of cDNA, and 3.5 μL of RNAse free water. The melting temperature was optimized for 55°C for all the primers tested. Primer gene sequences were designed using the NCBI primer (https://www.ncbi.nlm.nih.gov/tools/primer-blast/) tool and the Beacon designer software (http://www.premierbiosoft.com/molecular_beacons/) and can be found in **Supplementary Table S4**. The amplification efficiencies were tested for all primer pairs and were only used when efficiency is near 100%. To verify the specificity of the amplification and absence of primer dimer formation, corresponding melting curves were performed and analysed immediately after the amplification protocol. Non-specific products were not found in any case. *Oaz1* was used as endogenous control to normalize the expression levels of genes of interest and the relative mRNA expression levels were calculated using the delta C_T_ (2^-ΔΔCT^) method [35].

### 11. Statistical analysis

Statistical analysis was performed using GraphPad Prism (version 7.00, www. graphpad.com). For all datasets, tests for identification of outliers and normality were performed. To identify and remove likely outliers, the Robust Regression and Outlier Removal (ROUT) method with Q=10% was used. Following this, Gaussian distributions were tested using the Shapiro-Wilk normality test (α=0.05). When all datasets followed a normal distribution, statistical differences between groups were calculated based on one-way (influence of one factor) or two-way ANOVA (influence of two factors), followed by Dunnet’s or Tukey’s multiple comparisons test, respectively, for multiple comparisons. In case one dataset did not follow a normal distribution, non-parametric tests were performed. Statistical differences between groups were calculated based in Kruskall-Wallis analysis, followed by Dunn’s multiple comparisons test for multiple comparisons. A p-value below 0.05 was considered statistically significant and data are shown as mean ± standard deviation (SD).

## Results

### Oligodendrocyte differentiation can be recapitulated in 3D modified alginate hydrogels

We have previously reported the production of alginate hydrogels biofunctionalized with cell adhesive (RGD) and matrix metalloproteinase (MMP) peptides (PVGLIG) for optimal astrocyte growing and branching [21]. Through reductive amination we showed that peptide incorporation was dramatically increased allowing a more flexible adjustment of hydrogel formation parameters (such as gelation, stiffness, and degradability). Due to the opening of the polymer chain and peptide availability in oxidized alginates containing PVGLIG, astrocytic branching ability closely resembled the *in vivo* scenario.

Here, we sought to ascertain if alginate formulations containing PVGLIG and RGD were amenable to culture OLs and promote cell differentiation. RGD was chemically conjugated to high molecular weight (HMW) alginate through carbodiimide chemistry while PVGLIG was engrafted via carbodiimide chemistry or reductive amination on partially oxidized alginate (5 and 10% theoretical oxidation) [27, 36] (**Figure 1A** and **Supplementary Table S1**). Alginate hydrogels were formed by combination of alginate grafted with RGD and PVGLIG (**Figure 1B**) being the RGD concentration fixed to 40 µM. Previously we showed that the OL branching ability was increased for formulations with low RGD concentrations in comparison with higher concentrations (40 µM vs 100 µM) [21]. Here, we conducted a thorough process of OL culture optimization on alginate formulations with different oxidation and PVGLIG incorporation degrees. By varying the oxidation degree of the alginate chain, we expected an increasing on the mesh sizes with consequences for cellular behavior (higher mobility and increased space to branch out). Additionally, the variation of the concentration of the MMP-sensitive peptide would change its availability to the cells and therefore impact their capacity of remodel the 3D matrix.

We observed that all alginate hydrogels were soft with complex shear modulus below 1 kPa (**Figure 1C** and **Supplementary Table S2**), which are in the range of published brain’s mechanical properties [37]. While the presence of cells did not seem to influence the stiffness of the hydrogels (**Figure 1C**, accessed at day 1 of differentiation, D1 DIFF), the incorporation of PVGLIG either through carbodiimide or reductive amination significantly decreased the mechanical properties of the matrices. The most preeminent effects were observed for all formulations where PVGLIG was coupled by reductive amination (Alg Ox5 PVG 200 µM, Alg Ox10 PVG 200 µM, and Alg Ox10 PVG 400 µM, where “Ox” refers to oxidized alginate and is followed by a number representing the theoretical oxidation percentage). The decrease in stiffness was accompanied by an increase in cellular branching (**Figure 1D**), and metabolic activity (**Figure 1E**) for all time points tested (D5, 9, and 14 DIFF). No major effects were observed in terms of survival capacity (**Supplementary Figure 1A**). The expression of characteristic OL genes was also tendentially higher for oxidized formulations, particularly for the one with the highest concentration of PVGLIG (Alg Ox10 PVG 400 µM) (**Figure 1F**). The same formulation showed increased number of myelin basic protein (MBP) positive cells (**Supplementary Figure 1 B**) alongside with the lowest sphericity and the highest cell volume values (**Figure 1G, H** and **Supplementary Figure 1 C-F**). These findings are a strong indicative of enhanced OL differentiation capacity and for that reason we proceeded with the Alg Ox10 PVG 400 µM + RGD 40 µM for the subsequent studies presented in this work.

To test whether alginate hydrogels (Alg Ox10 PVG 400 µM + RGD 40 µM) were permissive not only for differentiation but also for OL utmost activity (wrapping), we cultured OLs on poly(dimethyl siloxane) (PDMS) micropillars,and embedded them with an alginate hydrogel (**Figure 1J**). Micropillars are cylindrical structures designed to function as axonal surrogates allowing myelination to occur in a biologically relevant platform. Based on previous conclusions regarding the myelination capacity of OLs in the different diameter micropillars, we used 5 µm pillars, with a height of 10 µm and interspaced 30 µm. For this setup an optimization of the micropillar surface characteristics was performed to maintain the alginate hydrogel intact throughout the cell culture period (**Supplementary Figure 2**). We consistently observed myelin rings around micropillars (**Figure 1J**), highlighting the relevance of our engineered alginate hydrogels in allowing myelination processes. To the best of our knowledge, this is the first time that a platform containing relevant three-dimensional features and axonal surrogates was described for the growing of OLs.

**Figure 1.**
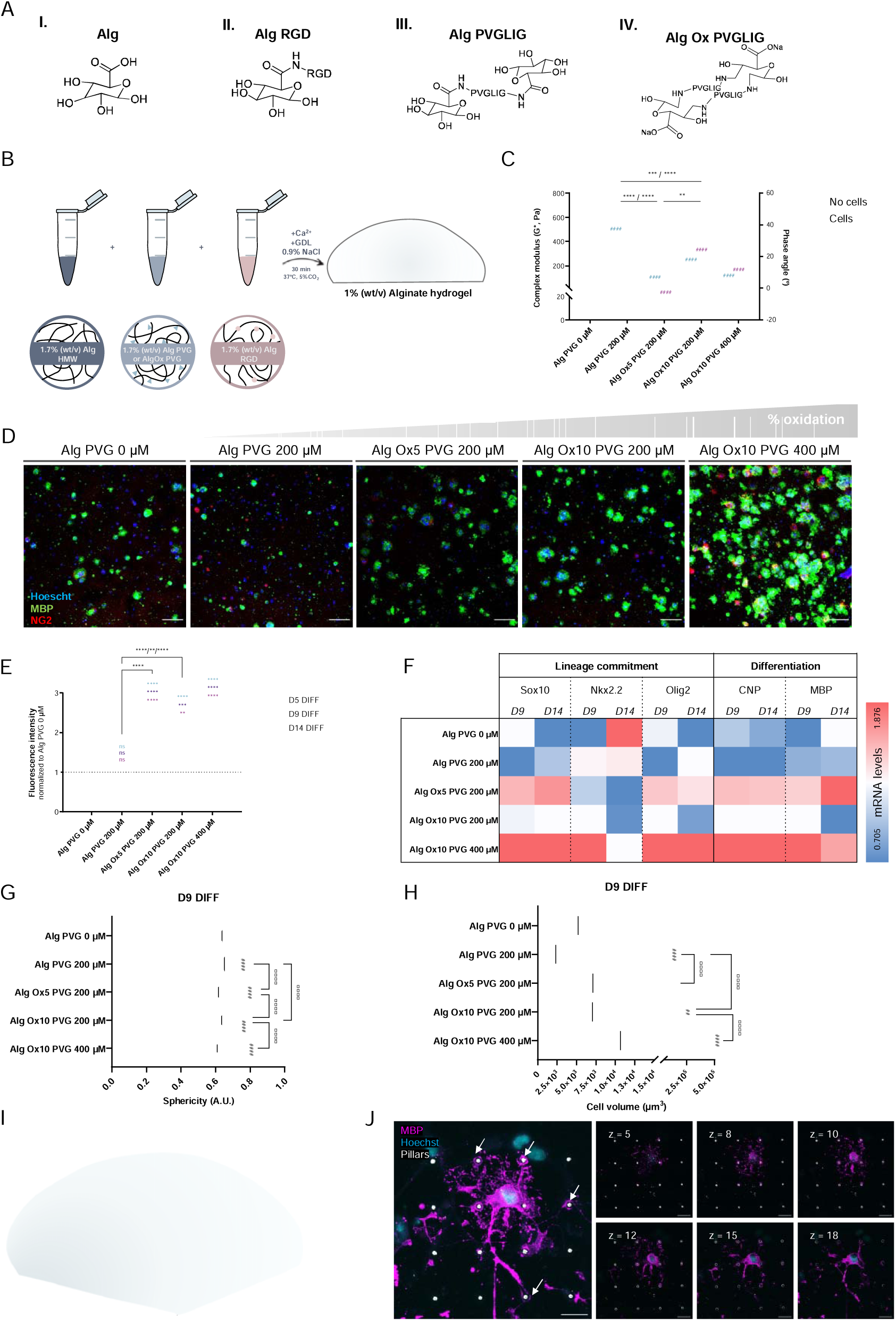
Alginate hydrogels containing PVGLIG and RGD are suitable matrices for the growing and differentiation of oligodendrocytes (OLs). **A.** Different alginate structures used in the present study. I – non-modified alginate, high molecular weight (HMW); II – alginate containing RGD coupled by carbodiimide chemistry; III – alginate containing PVGLIG coupled by carbodiimide chemistry; IV – alginate containing PVGLIG engrafted via reductive amination on partially oxidized alginate. **B.** Scheme of alginate hydrogel preparation. Cell-laden matrices were formed by mixing non-modified alginate (Alg HMW) with alginate containing PVGLIG and alginate containing RGD. RGD concentration was fixed at 40 µM and PVGLIG was varied to find the best alginate matrix for the growing of OLs. **C.** Rheological properties of alginate hydrogels estimated at day (D) 1 of differentiation (DIFF). n > 3 hydrogels per condition. Two-way ANOVA, Sidak’s multiple comparison test, cardinals represent statistical significance against Alg PVG 0 µM, #### p<0.0001. Asterisks represent statistical significance and “no cells” are listed before “cells”, **** p<0.0001, *** p<0.0002, ** p<0.0032, ns – not statistically significant. **D.** Representative confocal images of OLs in alginate hydrogels at D9 DIFF. Cells were labelled with myelin basic protein (MBP, green), Neuron-glial antigen 2 (NG2, red) and counterstained with Hoechst (blue). **E.** Metabolic activity profile of OLs in alginate hydrogels. Lines in the graphs represent mean values. n = 7-15 hydrogels per condition from 6 independent experiments. Two-way ANOVA, Tukey’s comparison test, asterisks represent statistical significance **** p<0.0001, ** p<0.0021, *p<0.032. Days of differentiation are listed in order in the graph. **F.** mRNA levels of OLs in alginate hydrogels at D9 and 14 DIFF. 2^-ΔΔCt^ values were normalized to the housekeeping gene *Oaz1*. n = 5-7 independent experiments (10-12 hydrogels pooled together per condition, per experiment). Statistical tests performed using One-way ANOVA with Tukey’s multiple comparison tests (independent tests for each individual gene and time-point). **G.** and **H.** Sphericity and cell volume (respectively) of OLs grown within alginate hydrogels at D9 DIFF. Sphericity values closer to 1 indicate a perfect sphere. n > 400 individual cells analyzed per condition. One-way ANOVA, Tukey’s multiple comparison test, cardinals represent statistical significance against Alg PVG 0 µM, #### p<0.0001. Asterisks represent statistical significance, **** p<0.0001. **I.** Scheme of the micropillars with OLs embedded within alginate hydrogels. **J.** Representative confocal image of OLs cultured in alginate hydrogels (Alg Ox10 PVG 400 µM) in the presence of poly(dimethyl siloxane) (PDMS) micropillars at D9 DIFF. Cells were labelled with MBP (magenta) and counterstained with Hoechst (cyan). Micropillars are in white. Different z planes are depicted indicating that cells can wrap the micropillars in a radial and transversal way when embedded within a three-dimensional matrix. Arrows are pointing to wrapped micropillars. Scale bar 25 µm.

### Stiffer and higher viscoelastic alginate matrices negatively impact the differentiation of oligodendrocytes along with alterations in Yap1/Taz expression

OLs are mechanosensitive cells [4] and mechanical properties are important regulators of nervous system development and disease. However, until now, 3D models to specifically study the impact of mechanical properties on these cells are scarce. After finding the optimal conditions for culturing OLs in alginate hydrogels (Alg Ox10 PVG 400 µM + RGD 40 µM) we tuned their mechanical properties and explored their impact on OL behavior and differentiation. By changing the polymer mass (1% wt/v, 2% wt/v, and 3% wt/v) as well as the ratio of oxidized to non-oxidized alginate (60:40, 30:70 and 20:80) we were able to progressively increase the complex shear modulus of the gels (**Figure 2A** and **Supplementary Table S3**) as well as their relaxation time (**Figure 2B**). These changes in mechanical properties were accompanied by a diminished cellular branching complexity and metabolic activity (**Figure 2C, D**), which was not translated into alterations in OL survival (**Supplementary Figure 3**). Additionally, we observed that the expression of characteristic OL genes was diminished for 2% Alg 30:70 and 3% Alg 20:80 in comparison with 1% Alg 60:40 (**Figure 2E**). Specifically, we noted that *Olig2* expression was significantly decreased for stiffer matrices at D9 and 14 DIFF, which is indicative of decreased commitment towards the OL lineage and arrest in progenitor states. Generally, we saw that the differentiation genes tested (*CNP* and *MBP*) were also decreased for cells cultured in those matrices, corroborating that apart from being less committed, these cells were also less differentiated. At a protein level, we quantified low number of cells expressing the myelin marker MBP for 3% Alg 20:80 matrices in comparison with 1% Alg 60:40 and 2% Alg 30:70 (**Figure 2F**). For 2% Alg 30:70 matrices, the number of MBP positive cells remained similar to the ones determined for the 1% Alg 60:40 one (**Figure 2F**) but the volume of the cells was tendentially diminished (**Figure 2G**), while no major changes were verified for sphericity (**Figure 2H**). These results reinforce that a 4-fold increase in the stiffness of hydrogels had a major impact on cellular morphology but not on myelin production, while a more dramatic change of the mechanical properties (17-fold increase) had pronounced effects on both, OL branching and differentiation. The difficulty of OLs in growing and differentiate might arise from the tighter mesh of the 3% Alg 20:80 alginate matrices (**Supplementary Table S3**).

Next, we investigated the most common mechanosensing routes that could be altered in OLs when cultured in stiffer and highly viscoelastic matrices. Assuming that the major differences on mechanosensing pathways would be seen at earlier time points due to the fast response of cells to mechanical cues [38], we evaluated OLs response at D1, 2 and 3 DIFF. Mechanotransduction pathways usually start in the cell periphery with receptors that connect the ECM to the cell’s cytoskeleton, as is the case of integrins. When external forces are applied, receptors undergo conformational changes, activating signaling cascades through proteins like focal adhesion kinase (FAK), also known as Ptk2. We evaluated the expression of *Ptk2* and one of its downstream targets, mitogen associated protein kinase 14 (*Mapk14*) also known as extracellular signal-regulated kinase (ERK), which are involved in focal adhesion kinase signaling pathways. A tendential decrease on the expression of both genes for the stiffer matrices was observed (**Supplementary Figure 4A, B**). Further, we found that at D2 DIFF the yes-associated protein 1 (Yap1) is mainly in the cytoplasm of OLs in 1% Alg 60:40 while it is found in the nuclei of OLs cultured in 2% Alg 30:70 and 3% Alg 20:80 (qualitative analysis) (**Figure 2I**). Moreover, we saw significant higher expression of *Yap1* and Tafazzin (*Taz*) genes in OLs cultured in 3% Alg 20:80 in comparison with 1 and 2% at D2 DIFF (**Figure 2J, K**). *Taz* expression for the stiffer hydrogels reached its highest expression at D2 DIFF, highlighting the importance of timing when analyzing mechanosensing events. Finally, mechanical forces transmitted through the cell cytoskeleton can also impact the nucleus and this can lead to alterations in chromatin remodeling. We also looked at Hdac1 expression profile, which is an enzyme involved in the regulation of genetic expression by modifying chromatin structure through removal of acetyl groups from histone proteins [39]. We also looked at possible epigenetic related genes (histone deacetylase 1, *Hdac1*) and saw a tendential increase in its expression at D2 DIFF for stiffer and high viscoelastic matrices (**Supplementary Figure 4C**).

**Figure 2.**
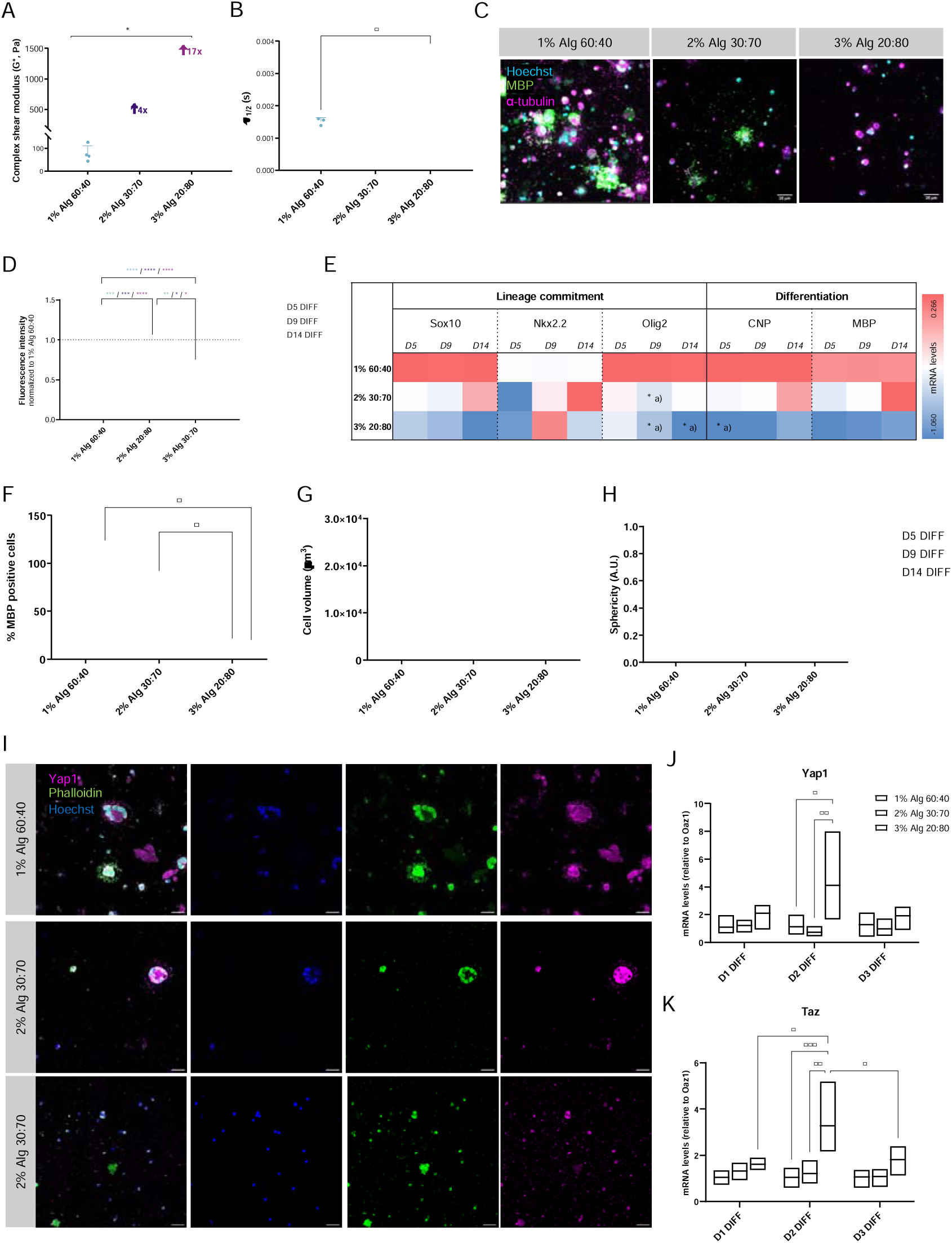
Alginate matrices with increased complex shear modulus and stress-relaxation times lead to decreased oligodendrocyte (OL) differentiation. **A.** Complex shear modulus of alginate matrices containing OLs at day 1 of differentiation (D1 DIFF). Alginate hydrogels were produced with increased polymer concentrations and different oxidized to non-oxidized ratios (heron designed 1% Alg 60:40, 2% Alg 30:70 and 3% Alg 20:80). n = 3-4 hydrogels per condition. Statistical tests were performed using One-way ANOVA, Kruskal-Wallis test, *p<0.05. **B.** Viscoelastic (stress-relaxation) properties of 1% Alg 60:40, 2% Alg 30:70 and 3% Alg 20:80 matrices at D1 DIFF (10% applied strain). n = 3-4 hydrogels per condition. Statistical tests were performed using One-way ANOVA, Kruskal-Wallis test, * p<0.032. **C.** Representative confocal images of OLs grown on the different matrices studied. Cells were labelled with myelin basic protein (MBP, green), α-tubulin (magenta) and counterstained with Hoechst (cyan). **D.** Metabolic activity of OLs in alginate matrices with different stiffness and viscoelasticity. Values were normalized to those obtained in 1% Alg 60:40 matrices. n = 5-9 hydrogels per condition from 4 independent experiments. Statistical tests were performed using One-way ANOVA, Kruskal-Wallis test, **** p<0.0001, *** p<0.0002, ** p<0.0021, * p<0.032. **E.** mRNA levels of OLs in different alginate matrices. Values were normalized to the housekeeping gene *Oaz1*. Colors indicate 2^-ΔΔCt^ mean values (minimum -1.060 and maximum 0.266). n = 3-5 independent experiments (total of 10-12 hydrogels pooled together *per* condition, per experiment). Statistics were performed using Two-way ANOVA, Tukey’s multiple comparison test. a) represents statistical differences against 1% Alg 60:40, ** p<0.0021, * p<0.032. **F. G.** and **H.** Quantification of the myelin basic protein (MBP) positive cells, cellular volume and sphericity at D5, 9 and 14 DIFF. n = 3 independent experiments. Statistics were performed using Two-way ANOVA, Tukey’s multiple comparison test, * p<0.032. **I.** Representative confocal images of OLs at D2 DIFF labelled with Yap1 (magenta) and Phalloidin (actin, green). Counterstaining was performed with Hoechst (blue). Scale bar represents 25 µm. **J.** and **K.** mRNA levels of Yap1 and Taz in OLs in alginate matrices at D1, 2 and 3 DIFF. 2^-ΔΔCt^ values were normalized to the housekeeping gene *Oaz1*. n = 3-4 independent experiments (total of 10-12 hydrogels pooled together per condition, per experiment). Statistics were performed using Two-way ANOVA, Tukey’s multiple comparison test, *** p<0.0002, ** p<0.0021, * p<0.032.

### Oligodendrocyte differentiation is impaired in alginate matrices with similar stiffnesses but higher viscoelastic properties along with alterations in Yap1/Taz, focal adhesion kinases and histone deacetylase expression

Stress-relaxation phenomena has gained crescent attention in the mechanobiology field, however, their comprehension in modulating CNS processes is still very poor. The brain is not a purely elastic material, instead as every other tissue in the human body, it presents a viscous component and exhibits a time-dependent response to cells traction forces [13, 14]. Matrix stress-relaxation properties describe the behavior of a material under constant strain or deformation over time. In a viscoelastic material the stress (force per area) decreases with time due to microstructural components rearrangements.

We next sought to understand the specific impact of the viscoelastic properties on OL differentiation. By increasing the polymer content in the hydrogels but at the same time maintaining the ratio of oxidized to non-oxidized alginate (60:40) we were able to produce matrices with similar stiffness (with complex shear moduli around 100 Pa) (**Figure 3A** and **Supplementary Table S2**) but with different stress-relaxation times (**Figure 3B**). We observed that OL branching decreased (**Figure 3C**) alongside with diminished metabolic activity (**Figure 3D**). However, cellular survival did not drop (**Supplementary figure S5**) for matrices with higher stress-relaxation values. In addition, OL differentiation characteristic genes were generally downregulated for these matrices (**Figure 3E**). Interestingly, *MBP* expression was increased for 2 and 3% Alg 60:40 at early time-points of differentiation (D5 DIFF) but as a function of the culture time its expression gradually decreased, indicating a possible failed attempt of OLs to sustain the higher expression of the myelin marker.

However, the increased expression of *MBP* was not translated in a higher number of cells positive for this marker (**Figure 3F**). The largest differences between high and low stress-relaxation matrices were seen at D9 and D14 DIFF, which are in accordance with the timing where MBP genetic expression lowered. The OL alterations were also observed in terms of cellular volume, with cells being smaller for 2 and 3% Alg 60:40 (**Figure 3G**) with no major differences in sphericity (**Figure 3H**).

Being mechanical properties the only factor changing among the different matrices we delve into the most common possible altered mechanotransduction pathways. By qualitative image analysis we did not observe a clear translocation of Yap1 to the nucleus in the presence of matrices with high stress-relaxation times (**Figure 3I**). Nevertheless, at a genetic level *Yap1* and *Taz* were augmented for the 3% Alg 60:40 hydrogels at all time-points tested (D1, 2, and 3 DIFF) which indicates a possible alteration in these co-transcriptional activators. We could also not discard that the selected time-point for evaluation of Yap protein translocation might not have been the most suitable one.

We observed a consistent decrease of the expression of *Ptk2* gene on OLs at D1, D2, and D3 DIFF for the 3% Alg 60:40 while for the 2% Alg 60:40 no major differences were seen (**Figure 3L**). Comparable to the expression pattern observed for *Ptk2*, we noted a significant decline in the expression of *Mapk14* for 2 and 3% Alg 60:40, particularly noticeable at D2 DIFF (**Figure 3M**). Mapk14 is involved in various cellular mechanisms including those related with stress responses, inflammation, cell differentiation, and apoptosis [40]. Specifically, its role in regulating cell fate decisions including differentiation is well-known. Thus, its decrease in matrices with high viscoelastic properties might be interconnected with impaired OL differentiation. Additionally, when looking at epigenetic regulators, we observed a massive increase on the expression of *Hdac1* for alginate matrices of 3% Alg 60:40 (**Figure 3O**). High *Hdac1* expression might impact the differentiation of OPCs into mature OLs. Changes in mechanical cues can trigger alterations in cellular differentiation, and Hdac1-mediated chromatin modifications might mediate these processes.

Overall, we proved that the culture of these cells in matrices with high complex shear modulus and stress-relaxation times led to alterations on the expression of mechano-associated transcription factors as *Yap1* and *Taz*, while focal adhesion proteins and chromatin remodeling events were not altered. On the other hand, in matrices with similar shear modulus but increased stress-relaxation times the impact on mechanosensing events was more pronounced with alterations on *FAK* signaling, *Yap1* and *Taz* transcription factors and chromatin remodeling *Hdac1*. This emphasizes the importance of independently comprehending the influence of viscoelasticity on neural cells. The tuning of this mechanical property could represent a new targeted approach in the framework of aging and neurological diseases.

Additionally, we also independently altered the complex shear modulus (varying from around 80 to 300 Pa) of alginate matrices without altering their viscoelastic properties (**Supplementary Figure S6 A, B**). We increased the polymer content from 1% to 2 and 3% (wt/v) by doubling and tripling the PVGLIG and RGD concentration. In this manner we maintained the ratio of oxidized to non-oxidized alginate similar (60:40). The metabolic activity of the cells, as well as the number of MBP positive cells drastically decreased only for the 3% alginate matrices while the cellular volume was diminished for both 2 and 3% alginate (**Supplementary Figure S6 C-G**). This also proves that stiffer matrices have a negative impact on OL differentiation independently of the viscoelastic properties, as already reported by others [41–43].

**Figure 3.**
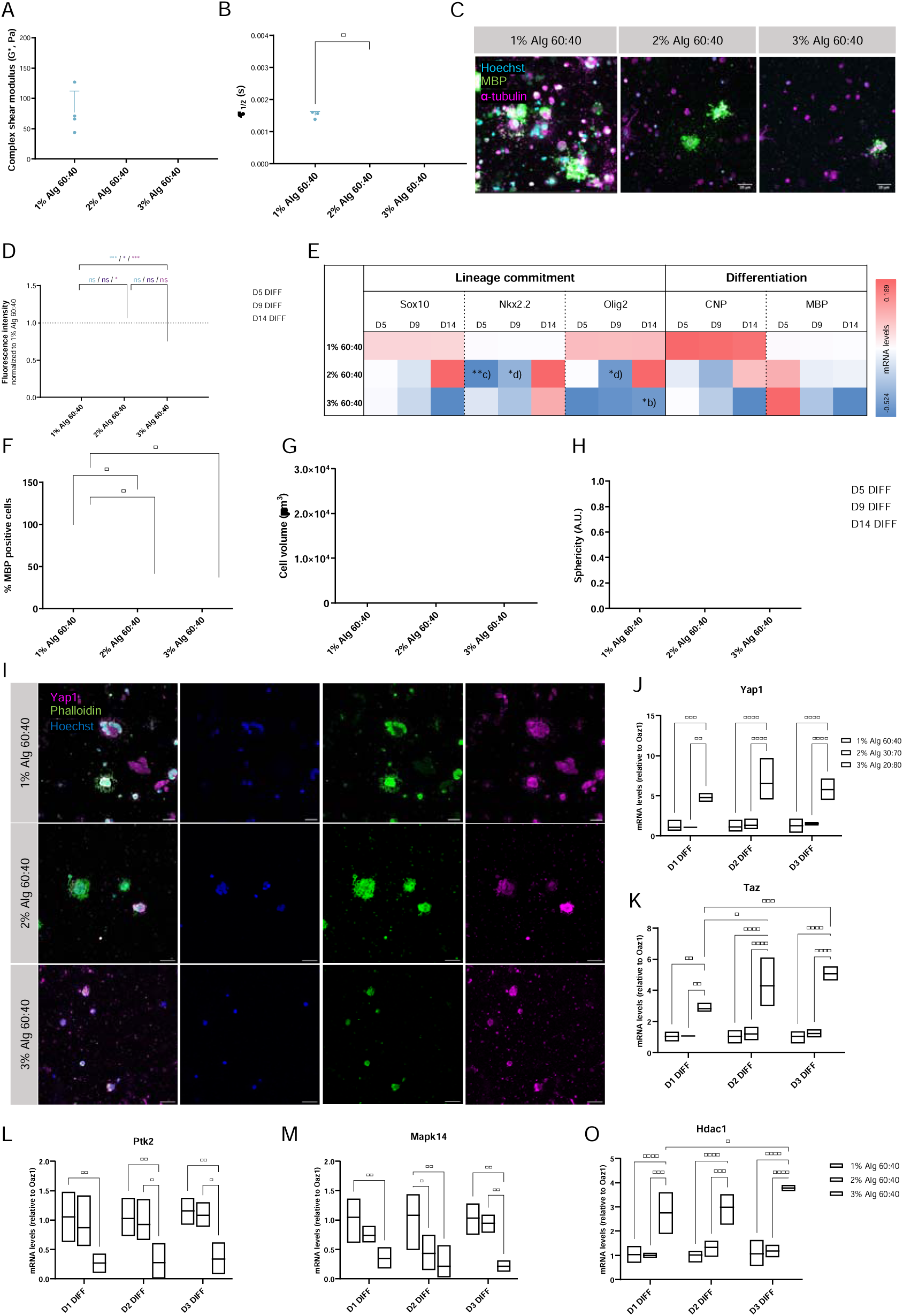
Alginate matrices with similar complex shear modulus but high stress-relaxation times lead to decreased oligodendrocyte (OL) differentiation. **A.** Complex shear modulus of alginate matrices containing OLs at day 1 of differentiation (D1 DIFF). Alginate hydrogels were produced with increased polymer concentrations and similar oxidized to non-oxidized ratios (heron designed 1% Alg 60:40, 2% Alg 60:30 and 3% Alg 60:40). n = 3-4 hydrogels per condition. Statistical tests were performed using One-way ANOVA, Kruskal-Wallis test, *p<0.05. **B.** Viscoelastic (stress-relaxation) properties of 1% Alg 60:40, 2% Alg 60:40 and 3% Alg 60:40 matrices at D1 DIFF (10% applied strain). n = 3-4 hydrogels per condition. Statistical tests were performed using One-way ANOVA, Kruskal-Wallis test, * p<0.032. **C.** Representative confocal images of OLs grown on the different matrices studied. Cells were labelled with myelin basic protein (MBP, green), α-tubulin (magenta) and counterstained with Hoechst (cyan). **D.** Metabolic activity of OLs in alginate matrices with similar stiffness but different viscoelasticity. Values were normalized to those obtained in 1% Alg 60:40 hydrogels. n = 5-9 hydrogels per condition from 4 independent experiments. Statistical tests were performed using One-way ANOVA, Kruskal-Wallis test, **** p<0.0001, *** p<0.0002, ** p<0.0021, * p<0.032. **E.** mRNA levels of OLs in different alginate matrices. Values were normalized to the housekeeping gene *Oaz1*. Colors indicate 2^-ΔΔCt^ mean values (minimum -0.524 and maximum 0.189). n = 3-5 independent experiments (total of 10-12 hydrogels pooled together per condition, per experiment). Statistics were performed using Two-way ANOVA, Tukey’s multiple comparison test. b) represents statistical differences against 2% Alg 60:40, c) represents differences between D5 and D14 DIFF and d) between D9 and D14 DIFF, ** p<0.0021, * p<0.032. **F. G.** and **H.** Quantification of the myelin basic protein (MBP) positive cells, cellular volume and sphericity at D5, 9 and 14 DIFF. n = 3 independent experiments. Statistics were performed using Two-way ANOVA, Tukey’s multiple comparison test, * p<0.032. **I.** Representative confocal images of OLs at D2 DIFF labelled with Yap1 (magenta) and Phalloidin (actin, green). Counterstaining was performed with Hoechst (blue). Scale bar represents 25 µm. **J. – O.** mRNA levels of Yap1, Taz, Ptk2, Mapk14 and Hdac1 in OLs in alginate matrices at D1, 2 and 3 DIFF. 2^-ΔΔCt^ values were normalized to the housekeeping gene *Oaz1*. n = 3-4 independent experiments (total of 10-12 hydrogels pooled together per condition, per experiment). Statistics were performed using Two-way ANOVA, Tukey’s multiple comparison test, *** p<0.0002, ** p<0.0021, * p<0.032.

## Discussion

Efficient signal propagation throughout neurons must be tightly coordinated and organized to allow us to perform our daily life activities. Myelin, the membranous sheath that surrounds axons and is responsible for increasing the velocity of signal propagation, evolved in a way that allowed vertebrates to achieve functional nervous systems. The significance of myelin to the nervous system is being increasingly recognized and emerging as a potential key factor in regulating developmental [44, 45], adult [46], and age-related [47] disorders of the CNS. Formation of the myelin sheath, wrapping and growing in length and thickness involve an intimate alignment of many factors including timing, precise location, and molecular and biophysical cues from the axons and the surrounding environment [4, 48]. Specifically, research on the relevance of the mechanical properties of the brain has demonstrated that physical factors not only contribute to disease progression but also can also be their cause [49]. How do mechanical cues dictate OL differentiation and myelination? Can we use this knowledge to model OL behavior and create new relevant therapies?

Some studies have delved into the mechanical environment influence on OLs. Seminal works using 2D *in vitro* substrates with different stiffnesses have concluded that OLs are mechanosensitive and that the ideal mechanical properties change depending on their stage of differentiation [6, 7, 9, 12, 41, 42, 50]. Nonetheless, the majority of the published works are based on cells growing on flat surfaces in two-dimensions which poorly mimic the *in vivo* scenario. Hence, it urges a need for the development of 3D models that can recapitulate OL differentiation and at the same time allow the specific study of mechanical properties effects on OL biology. Such models would be valuable not only to study fundamental cellular mechanisms but also to find novel therapeutic targets and serve as drug testing platforms.

Here we aimed to develop 3D hydrogel-based matrices amenable for culturing OLs and deepening the understanding of brain’s mechanical properties impact on the course of oligodendroglial cells behavior. Based on our former works with astrocytes [21, 23] we chose alginate as our starting 3D backbone matrix due to (1) its tunable mechanical properties; (2) the easiness of production and modification with cell-relevant moieties; and (3) the possibility of retrieving cells for further cellular and molecular biology studies.

Previously, we showed that the incorporation of the cell-adhesive RGD, and the MMP-sensitive PVGLIG peptides was advantageous for the growing and branching out of astrocytes and that the chemical route chosen for grafting the peptides had a tremendous impact on cellular behavior. Specifically, oxidizing alginate to 5% theoretical ratio led to increased grafting of PVGLIG comparing with modifying alginate in its pure state [21]. Here, as OLs are also MMP producers (specifically MMP-9) [51], we used the same rationale to create a biologically relevant matrix to culture OLs. By playing with oxidation molar ratios and with PVGLIG concentrations, we observed that the OL metabolic activity, branching out, and differentiation pattern were boosted in matrices with high PVGLIG concentrations (400 µM) and high oxidation (10%). Although we found that OLs express high levels of myelin related markers for the formulation containing 5% oxidation (Alg Ox5 PVG 200 µM), its extreme low complex shear modulus made these matrices highly fragile and susceptible to damaging. The reduced stiffness relative to matrices with similar PVGLIG concentration but higher oxidation levels, is explained by the low incorporation of the peptide in Alg Ox5. The lower percentage of oxidation compared to Alg Ox10 results in fewer reactive sites, leading to less peptide engrafted. Compensation for this lower peptide integration requires the addition of more oxidized alginate to achieve comparable final concentrations of peptide. Overall, our findings validated prior results regarding incorporation of PVGLIG via carbodiimide chemistry versus reductive amination: cells acquire a more relevant phenotype when choosing the latter chemical route. The oxidation of the alginate opens the polymer chain, increasing its interspace and allowing cells to move more freely while extending processes. Non-oxidized alginates with more compacted polymer chains have tighter mesh sizes, which can hinder OLs’ process elongation and subsequently their differentiation capacity.

Importantly, we proved that in these matrices, OLs can effectively perform their principal function: myelination. When the optimal hydrogel formulation (Alg Ox10 PVG 400 µM + 40 µM) was put in contact with a PDMS micropillar array previously described by us, we observed that after 9 days of growing in differentiating conditions, OLs were able to wrap the structures not only radially but also transversally. Up until now, only one model documented in the literature has incorporated electrospun fibers to culture Schwann cells, embedded within a fibrin hydrogel, aiming to mimic peripheral nerves [52]. In this study, we report, for the first time, a 3D platform specifically engineered to be permissive for CNS myelination. Our design incorporates transparent micropillars arranged uniformly in an upright position, enabling straightforward assessment of the OL wrapping ability.

Taking advantage of the tunable mechanical properties of alginate we proved that not only stiffness but also the viscoelasticity of the 3D matrix is fundamental in modelling OL behavior, a parameter that has been neglected for long. Our brain is not only ultra-soft, but also contains a high water content (approximately 80%) in comparison with other soft tissues in the human body, which makes it an almost perfect viscoelastic material [53]. When subjected to mechanical forces the brain can display elastic and viscoelastic behaviors: it can deform and return to its original shape or after deformation, it dissipates energy and has time-dependent behavior. All depends on the context. Here we showed that matrices with shear modulus around 1.3 kPa and high-stress relaxation times negatively impact the course of OL differentiation without major effects on cell viability. When increases of stiffness were made at a lower extent (from 0.1 to 0.3 kPa), stress-relaxation properties were maintained, and similarly this led to decreased OL differentiation. Finally, when we maintained the stiffness of the hydrogels around 0.1 kPa and increased the stress-relaxation properties we saw again diminished OL branching and differentiation capacities. Altogether our results prove that increasing stiffness or viscoelasticity whether independently or in combination, has a negative impact on the OL differentiation. These findings are in line with previous *in vitro* [5, 43, 50] and *in vivo* studies where stiffening of the matrix as a consequence of chronic demyelination decreased the capacity of OL myelination [54]. Although there are no published studies on the specific impact of viscoelasticity on neural cells, research with mesenchymal stem cells proved that parameters as cell spreading and proliferation are improved on matrices with low stress-relaxation times [15]. This corroborates the increased branching capacity of OLs on matrices with faster relaxation. We hypothesized that cells within these matrices can dissipate energy in a more efficient way than in slower relaxation ones. This allows the continuous generation of more energy that will be translated in process extension and branching-out.

Concomitantly with OL phenotypic changes we also observed alterations in the most common mechanosensing pathways. When stiffness and viscoelasticity were together augmented, gene expression data revealed alterations mainly at the transcriptional factors *Yap1* and *Taz*. Those mechanosensors were pointed by others as mediators of OL differentiation in fibers with tunable stiffnesses [5] and were also altered when OLs were mechanically stretched [55].

More surprisingly we found that viscoelastic alterations independently of stiffness changes alter the expression of many other mechanotransduction genes. Apart from *Yap1* and *Taz*, OLs in matrices with higher stress-relaxation times showed decreased expression of genes involved in FAK pathway (*Ptk2* and *Mapk14*) which were already proven to be intimately connected to the regulation of OL differentiation [56, 57]. A decrease in *Ptk2* expression might imply alterations in the formation of adhesion complexes affecting how cells adhere and interact with matrices. Although the concentration of the cell-adhesive peptide RGD was maintained constant in the different hydrogels, the denser mesh of slow relaxation matrices could have hindered peptide presentation to the cells. Therefore, we hypothesized that the adhesion of OLs to these matrices is challenged due to the limited capacity of the cell-binding domains to move in the alginate mesh and bind to the cellular integrins. The same trend was observed in matrices with increased stiffness and stress-relaxation times, but at lower extent. This suggests that cells have increased capacity of adapting in viscoelastic matrices comparing with elastic ones, showing diminished cellular responses in the latter. Nonetheless, further studies to confirm matrix remodeling by OLs in different matrices should be conducted (for example, the assessment of MMP-9 production). We also found alterations on the OLs at the level of the nuclear mechanotransduction (*Hdac1* expression), which were not observed when these cells were cultured in matrices with different stiffness and viscoelasticity properties. This suggests either the existence of a different regulatory mechanism behind OL behavior depending on the mechanical cue or once again an exacerbated response elicit in cells by viscoelastic matrices due to their capacity of adapting to the matrix overtime. An ongoing in-depth analysis of the differentially expressed genes in these matrices is being conducted (RNA sequencing analysis).

Many disease models only consider the elastic component of the brain [13], however the viscoelastic properties should not be neglected as they are reported to vary with factors as development, aging, and in disease context [20]. Although the extension of viscoelastic changes is still poorly understood, evidence points to an increased stress-relaxation response during aging in rodent models [20], an indicative of a decreased capacity of dissipating energy and remodel the matrix. This might be one of the explanations of the limited capacity of aged brains in resolving tissue degeneration. Still, the implications of dynamic changes in viscoelasticity for brain function and vulnerability need to be further investigated. Here we give strong evidence that viscoelasticity is a crucial player in modulating glial cell responses, with slower relaxation matrices leading to decreased OL differentiation. Based on these findings we hypothesized that viscoelasticity could represent a new target approach to modulate oligodendroglial behavior over disease and aging course. Here we point alterations in adhesion kinases and epigenetic histone deacetylases as potential biological targets to model how OLs feel viscoelasticity changes.

Taking into the consideration the lack of suitable models designed to have tunable stress-relaxation times, our hydrogels constitute an important advance for the current gap in the understanding of viscoelastic dimension to the brain function [15, 16].

Research on the mechanobiology of OLs can provide insights into disease mechanisms, aid in diagnosis, and potentially guide the development of new therapeutic approaches.

Our easy, versatile, 3D, and brain compliant platform could further help in the study of mechanisms regulating myelination processes and open further avenues in the search for biological targets promoting OL differentiation.

