## Supplementary Information for "Matrix stress-relaxation and stiffness modulate oligodendrocyte differentiation"

**Matrix stress-relaxation and stiffness modulate oligodendrocyte differentiation: an alginate-based approach**

Eva D. Carvalho^1,2,3^, Miguel R. G. Morais^1,2,4^, Georgia Athanasopoulou^1,2,4^, Marco Araújo^1,2^, Hendrik Hubbe^5^, Sofia C. Guimaraes^1,2^, Eduardo Mendes^5^, Stefano Pluchino^6^, Cristina C. Barrias^1,2,4^, Ana P. Pêgo^1,2,4*^

\# equal contribution

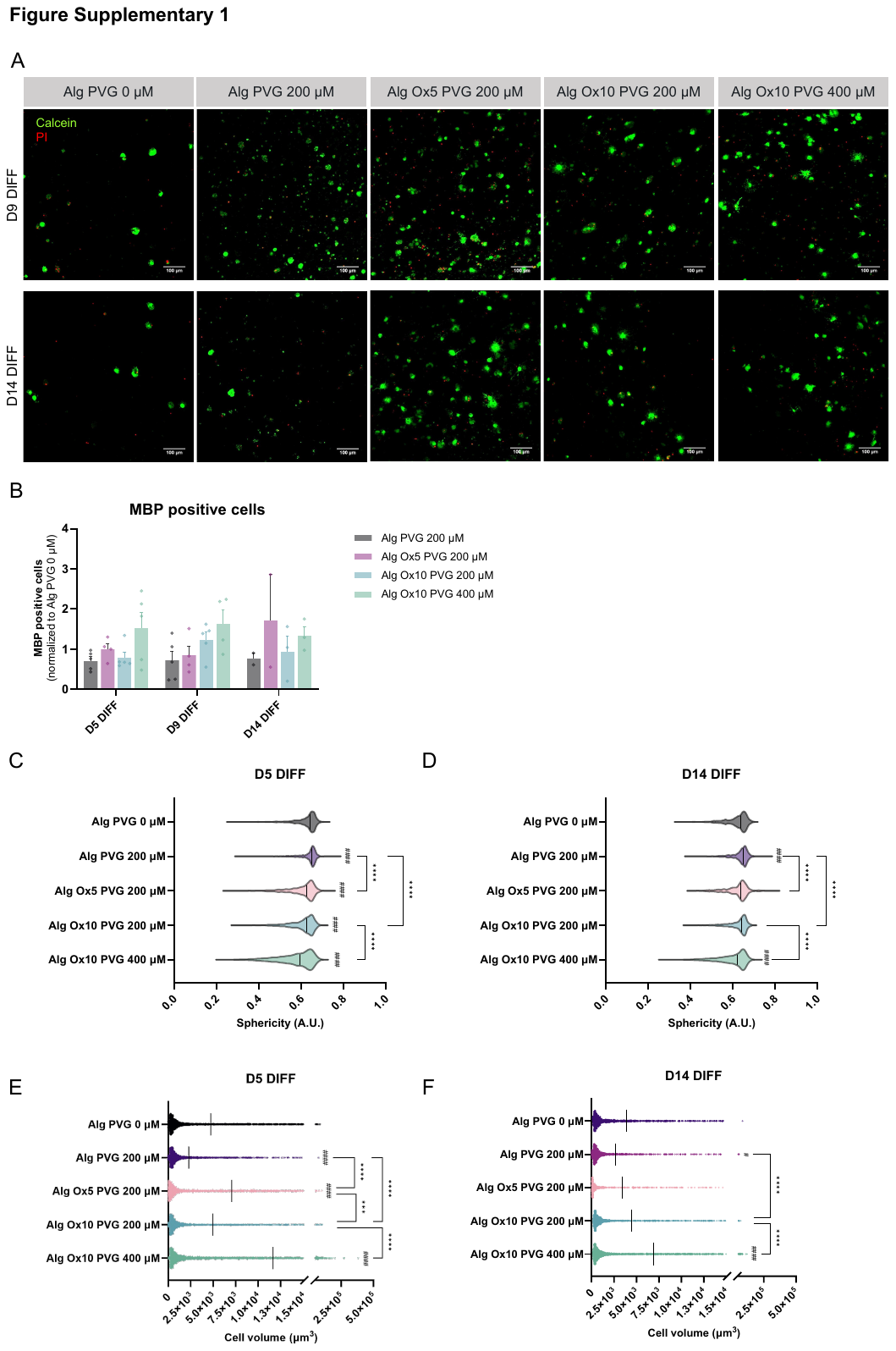

**Figure S1** Oligodendrocyte (OL) viability and differentiation in alginate hydrogels. **A.** Representative images of OL viability in alginate hydrogels at D9 and 14 of cell differentiation (DIFF). OLs were stained with Calcein-AM (green, showing the live cells) and propidium iodide (PI, red, representing the death cells). Scale bar 100 µm. **B.** Quantification of myelin basic protein (MBP) positive cells for OLs in alginate hydrogels at D5, D9 and D14 DIFF. n = 2-5 alginate hydrogels from 2-4 independent experiments. **C.** and **D.** Quantification of cell sphericity at D5 and D14 DIFF, respectively. Sphericity values closer to 1 indicate a perfect sphere. n > 400 individual cells analyzed per condition. One-way ANOVA, Tukey’s multiple comparison test, cardinals represent statistical significance against Alg PVG 0 µM, #### p<0.0001. Asterisks represent statistical significance, **** p<0.0001. **E.** and **F.** Cell volume of OLs grown within alginate hydrogels at D5 and D9 DIFF, respectively. n > 400 individual cells analyzed per condition. One-way ANOVA, Tukey’s multiple comparison test, cardinals represent statistical significance against Alg PVG 0 µM, #### p<0.0001. Asterisks represent statistical significance, **** p<0.0001, *** p<0.0002, * p<0.032.

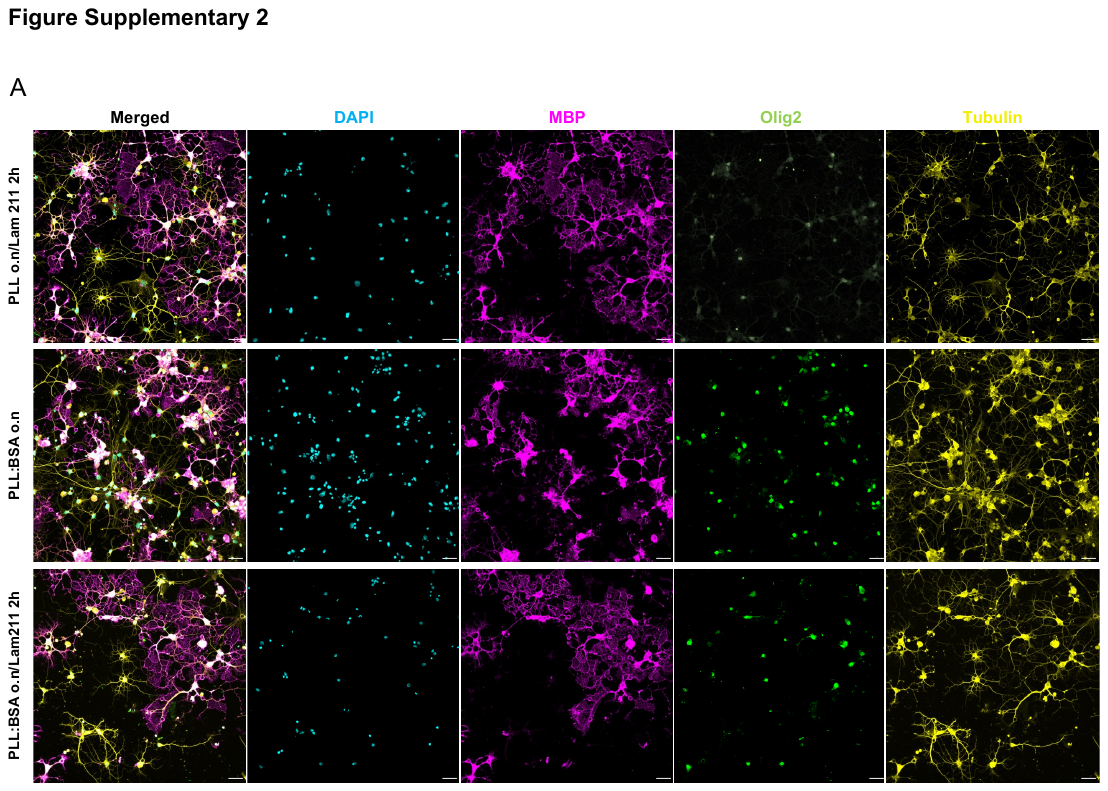

**Figure S2** Morphology of oligodendrocytes (OLs) cultured on micropillars with different coatings at day 5 of differentiation. No visible differences were observed on OL morphology and differentiation pattern after coating with poly-L-lysine (PLL) and bovine serum albumin (BSA) mixture overnight either or not followed by laminin 211 coating. Scale bar 25 µm.

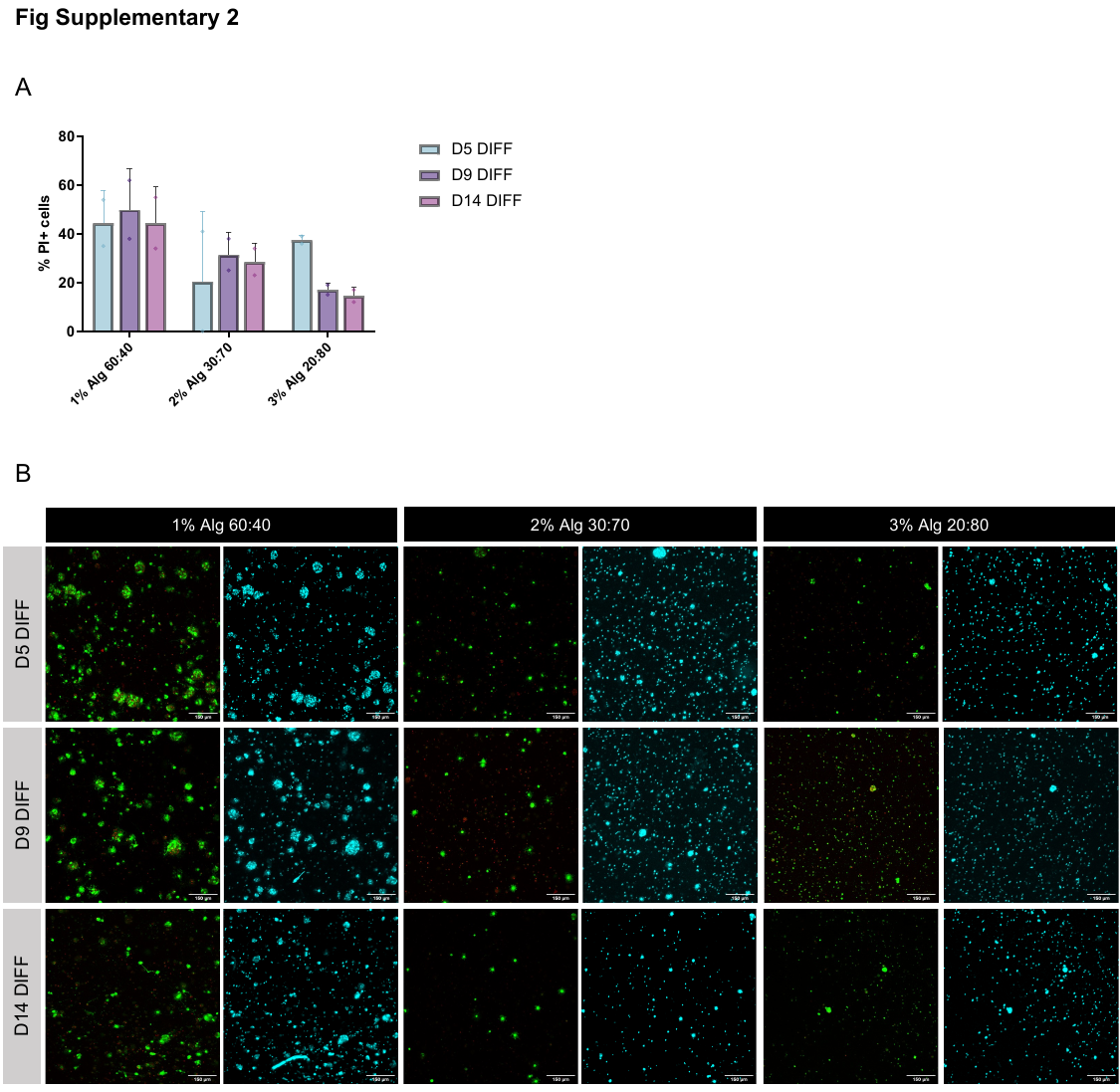

**Figure S3** Oligodendrocyte (OL) viability in alginate hydrogels of crescent shear modulus and stress-relaxation. **A.** Quantification of the number of propidium iodide positive cells at day 5, 9 and 14 of differentiation (DIFF). n = 2 hydrogels per condition **B.** Representative images of OL viability in alginate hydrogels at D5, 9 and 14 DIFF. OLs were stained with Calcein-AM (green, showing the live cells) and propidium iodide (PI, red, representing the death cells). Scale bar 100 µm. 1%, 2% and 3% indicate the percentage of alginate in the hydrogels. 60:40, 30:70 and 20:80 represent the ratio of oxidized to non-oxidized alginate.

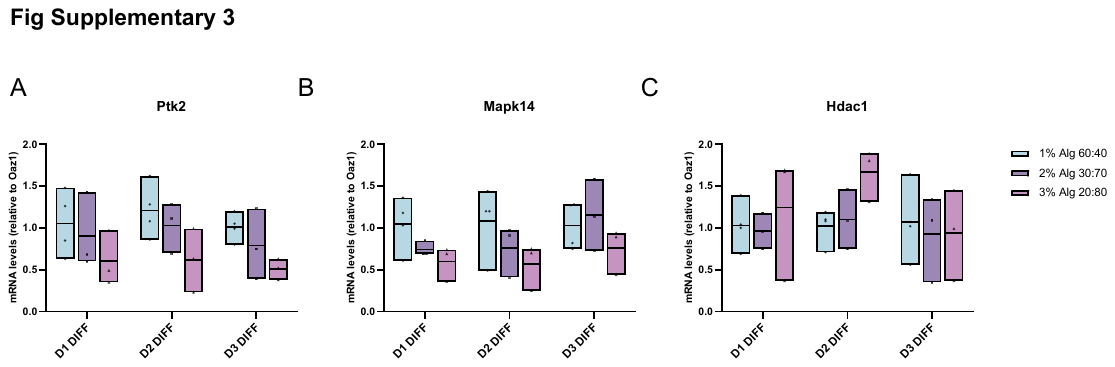

**Figure S4** mRNA levels of Ptk2, Mapk14 and Hdac1 (**A**, **B**, **C**) in oligodendrocytes in alginate matrices at D1, 2 and 3 DIFF. 2^-ΔΔCt^ values were normalized to the housekeeping gene *Oaz1*. n = 3-4 independent experiments (total of 10-12 hydrogels pooled together per condition, per experiment). Statistics were performed using Two-way ANOVA, Tukey’s multiple comparison test.

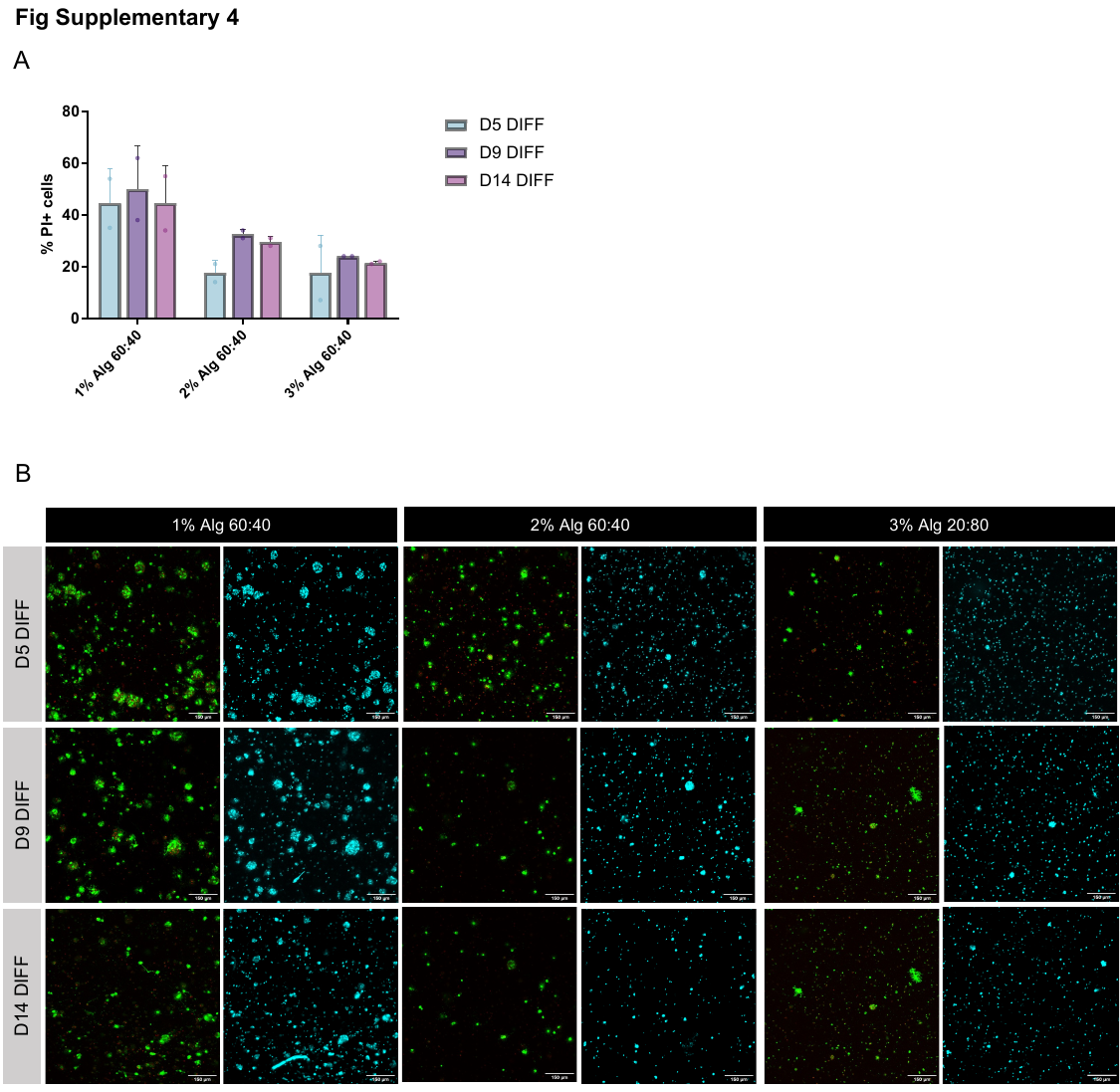

**Figure S5** Oligodendrocyte (OL) viability in alginate hydrogels of crescent stress-relaxation and similar shear modulus. **A.** Quantification of the number of propidium iodide positive cells at day 5, 9 and 14 of differentiation (DIFF). n = 2 hydrogels per condition **B.** Representative images of OL viability in alginate hydrogels at D5, 9 and 14 DIFF. OLs were stained with Calcein-AM (green, showing the live cells) and propidium iodide (PI, red, representing the death cells). Scale bar 100 µm. 1%, 2% and 3% indicate the percentage of alginate in the hydrogels. 60:40 represents the ratio of oxidized to non-oxidized alginate.

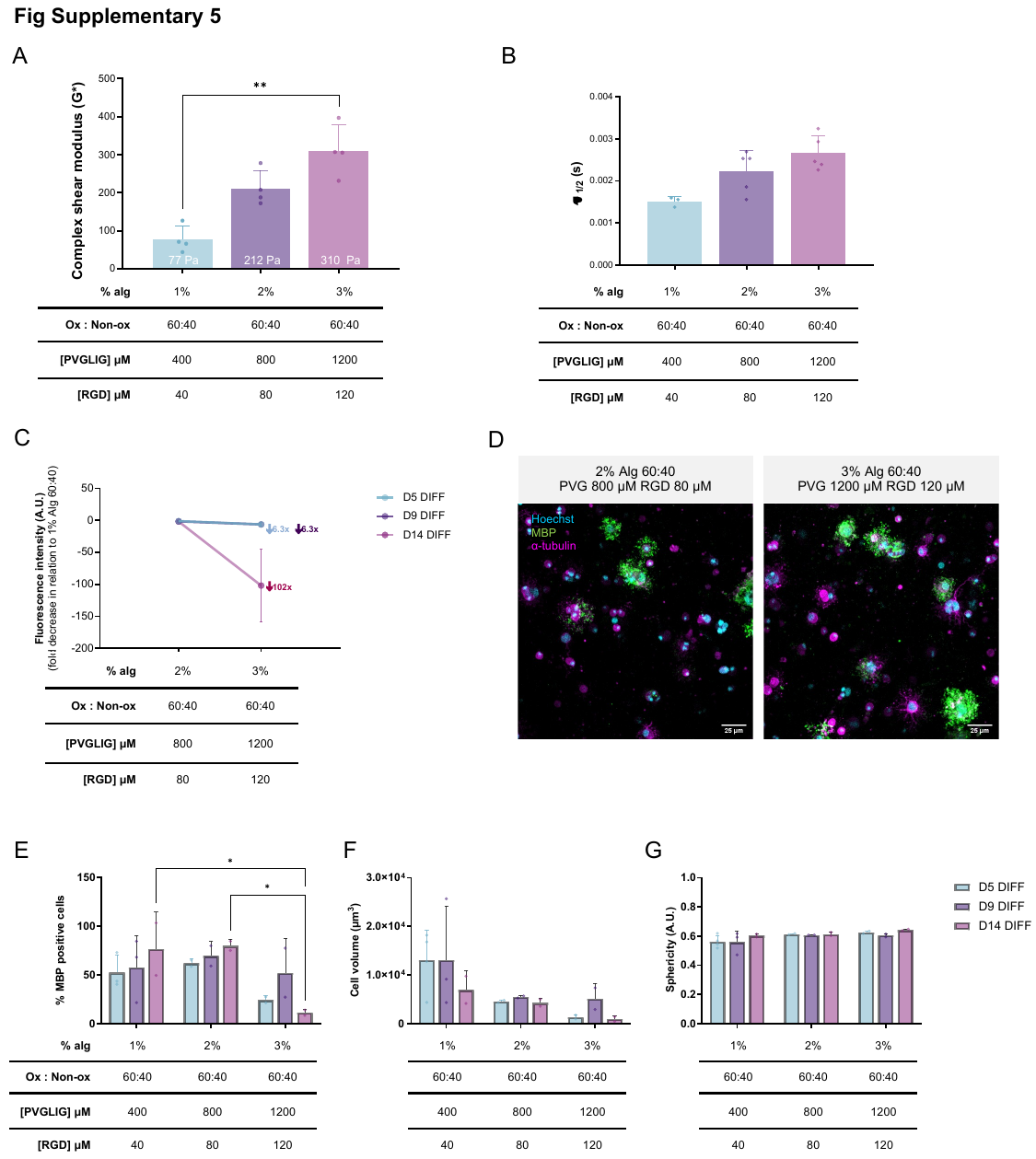

**Figure S6** Oligodendrocyte (OL) behavior in matrices with increased complex shear modulus and similar stress-relaxation times. **A.** Complex shear modulus of alginate matrices containing OLs at day 1 of differentiation (D1 DIFF). n = 2-4 hydrogels per condition. Statistical tests were performed using One-way ANOVA, Kruskal-Wallis test, *p<0.05. Alginate hydrogels were produced with increased polymer concentrations, similar oxidized to non-oxidized ratios and increased concentration of RGD and PVGLIG (40, 80 and 120 µM RGD and 400, 800 and 1200 µM PVGLIG. **B.** Viscoelastic (stress-relaxation) properties of matrices at D1 DIFF (10% applied strain). n = 3-4 hydrogels per condition. Statistical tests were performed using One-way ANOVA, Kruskal-Wallis test. **C.** Metabolic activity of OLs in alginate matrices with different stiffness and viscoelasticity. Values were normalized to those obtained in 1% Alg 60:40 matrices. Statistical tests were performed using One-way ANOVA, Kruskal-Wallis test. **D.** Representative confocal images of OLs grown on the different matrices studied. Scale bar 25 µM. **E.**, **F.**, and **G.** Quantification of the myelin basic protein (MBP) positive cells, cellular volume and sphericity at D5, 9 and 14 DIFF. n = 2-3 independent experiments. Statistics were performed using Two-way ANOVA, Tukey’s multiple comparison test, * p<0.032.

**Supplementary Table S1** List of alginate oxidized at a theoretical ratio of 10% with PVGLIG incorporated (theoretical peptide incorporation of 70 mg/g of alginate). The oxidation was estimated based on the percentage of non-reacted aldehydes via tert-butyl carbazate synthesis. Peptide incorporation was calculated using the DC kit (Bio-Rad). “B” stands for “batch” and “R” for “reaction”.

| **Batch number** | **Batch ID** | **% real oxidation** | **Peptide incorporation (mg/g)** |
| --- | --- | --- | --- |
| 1 | Alg Ox10 B2 PVG R1 | 16.91 ± 0.52 | 69,30 |
| 2 | Alg Ox10 B3 PVG R1 | 16.60 ± 0.70 | 66,28 |
| 3 | Alg Ox10 B4 PVG R2 | 18.17 ± 1.95 | 68,67 |
| 4 | Alg Ox10 B6 PVG R1 | 14.80 ± 1.05 | 61,81 |
| 5 | Alg Ox10 B5 PVG R2 | 13.33 ± 1.24 | 71,21 |
| 6 | Alg Ox10 B5 PVG R3 | 13.33 ± 1.24 | 64,03 |
| 7 | Alg Ox10 B6 PVG R2 | 14.80 ± 1.05 | 62,99 |
| 8 | Alg Ox10 B6 PVG R4 | 14.80 ± 1.05 | 73,94 |
| 9 | Alg Ox10 B7 PVG R1 | 14.8 | 64,97 |
| 10 | Alg Ox10 B7 PVG R2 | 14.8 | 70,12 |
| 11 | Alg Ox10 B7 PVG R3 | 14.8 | 75,05 |
| 12 | Alg Ox10 B7 PVG R4 | 14.8 | 68,04 |

**Supplementary Table S2** Rheological properties of alginate hydrogels with PVGLIG engrafted via carbodiimide chemistry or reductive amination (including “Ox” in the sample identification) containing different oxidation degree and PVGLIG concentrations. G* is the complex shear modulus, G’ is the elastic shear modulus, G’’ is the viscous shear modulus and 𝜉 is the mesh size.

| **Sample** | **Presence of cells** | **G* (Pa)** | **G' (Pa)** | **G'' (Pa)** | **Phase angle (º)** | **𝜉 (nm)** |
| --- | --- | --- | --- | --- | --- | --- |
| Alg PVG 0 μM | No | 559,48 ± 41,28 | 558,6 ± 41,00 | 25,56 ± 3,75 | 2,26 ± 0.47 | 19,73 ± 0,47 |
|  | Yes | 496,19 ± 41,41 | 482,71 ± 43,96 | 112,19 ± 6,84 | 13,20 ± 1,99 | 20,72 ± 0,64 |
| Alg PVG 200 μM | No | 340,65 ± 83,87 | 287,15 ± 33,98 | 148,07 ± 113,74 | 23,84 ± 13,87 | 24,39 ± 0,92 |
|  | Yes | 387,30 ± 70,79 | 265,24 ± 34,68 | 289,20 ± 68,09 | 46,01 ± 3,96 | 25,34 ± 1,12 |
| Alg Ox5 PVG 200 μM | No |  |  |  |  |  |
|  | Yes |  |  |  |  |  |
| Alg Ox10 PVG 200 μM | No | 159,07 ± 60,16 | 135,01 ± 53,97 | 83,78 ± 27,34 | 32,32 ± 2,18 | 32,34 ± 3,91 |
|  | Yes | 187,40 ± 30,35 | 178,80 ± 33,94 | 53,85 ± 9,49 | 17,34 ± 5,73 | 28,99 ± 1,98 |
| Alg Ox10 PVG 400 μM | No | 65,07 ± 22,09 | 62,08 ± 19,26 | 15,48 ± 17,22 | 9,95 ± 13,23 | 41,71 ± 5,01 |
|  | Yes | 76,86 ± 42,93 | 56,44 ± 17,16 | 46,96 ± 50,10 | 35,12 ± 21,64 | 42,88 ± 5,17 |

**Supplementary Table S3** Rheological properties of alginate hydrogels with different stiffness and stress-relaxation properties. G* is the complex shear modulus, G’ is the elastic shear modulus, G’’ is the viscous shear modulus and 𝜉 is the mesh size. 60:40, 30:70 and 20:80 represent the ratio of oxidized to non-oxidized alginate.

| **Sample** | **G* (Pa)** | **G' (Pa)** | **G'' (Pa)** | **Phase angle (º)** | **𝜉 (nm)** |
| --- | --- | --- | --- | --- | --- |
| 1% Alg 60:40 PVG 400 μM RGD 40 μM | 76,86 ± | 54,44 ± | 46,96 | 35,12 | 42,88 |
| 2% Alg 30:70 PVG 400 μM RGD 40 μM | 325,02 | 323,56 | 30,78 | 5,28 | 23,82 |
| 3% Alg 20:80 PVG 400 μM RGD 40 μM | 1304,73 | 1297,3 | 138,83 | 6,09 | 14,89 |
| 2% Alg 60:40 PVG 400 μM RGD 40 μM | 109,35 | 108,71 | 10,48 | 5,53 | 34,57 |
| 3% Alg 60:40 PVG 400 μM RGD 40 μM | 121,58 | 120,97 | 12,07 | 5,7 | 33,85 |
| 2% Alg 60:40 PVG 800 μM RGD 80 μM | 211,56 | 211,13 | 11,12 | 2,83 | 27,47 |
| 3% Alg 60:40 PVG 1200 μM RGD 120 μM | 351,18 | 335,18 | 26,75 | 4,56 | 24,21 |

**Supplementary Table S4** Primer sequences used for oligodendrocyte gene expression studies. Forward and reverse sequences are shown.

| **Primer name** | **Forward sequence (5'-3')** | **Reverse sequence(5'-3')** |
| --- | --- | --- |
| **MBP** | TGTCACAATGTTCTTGAAGAA | GCTCCCTGCCCCAGAAGT |
| **CNP** | GCTTCGACACTTCATTTCTG | GTCTCTTGCCAAAATAGCTG |
| **Nkx2.2** | TGCCCCTTAAGAGTCCTTT | TCCGTGCAGGGAGTATT |
| **Sox10** | AAGCTCTGGAGGTTGCTG | CGAGGTTGGTACTTGTAGTC |
| **Olig2** | TGGGTGTCAGAAACACTTAG | CGTTAGGAAACCACAAATCG |
| **Yap1** | TGGTGAGAGGTAGCAGAG | GCAAGATGAGAGCGAAGT |
| **Taz1** | AAGGAAGTGCTGTATGAG | AAGACTGGTGGTTAGAGA |
| **Hdac1** | GAGCAAGATGGCGCA | GCTTTGTGAGGACGATAGA |
| **Ptk2** | GGAGTCATTACAGAGAAC | GCTGATAGGCATATAAGAT |
| **Mapk14** | GATTATGCTGAATTGGAT | TGGTTAATATGGTCTGTA |
| **Oaz1** | TGCAGCAGCGAGAGTTCTAGG | CCGGACCCAGGTTACTACAG |

*MBP – myelin basic protein, CNP – 2',3'-Cyclic nucleotide 3'-phosphodiesterase, Nkx2.2 – homeodomain protein NK2 homeobox 2, Sox10 – SRY-related HMG-box 10, Olig2 – oligodendrocyte transcription factor 2, Yap1 – yes-associated protein 1, Taz – Phospholipid-Lysophospholipid Transacylase, Hdac 1 – histone deacetylase , Ptk2 – focal adhesion-associated protein kinase 2, Mapk14 – mitogen-activated protein kinase 14, Oaz1 – ornithine decarboxylase antizyme 1.*

// Macro "Break up a lif into individual TIFF"

// Open the file manager to select a lif file to break it into TIFFs

// In this case, only the metadata specific to a series will be written

// This macro also splits channels from confocal SP8 images

// Author: Mafalda Sousa, ALM platform, i3S

// 2020

path = File.openDialog("Select a File");

out_path = getDirectory("Choose Destination Directory ");

print(out_path);

run("Bio-Formats Macro Extensions");

Ext.setId(path);

Ext.getCurrentFile(file);

Ext.getSeriesCount(seriesCount);

setBatchMode(true);

for (s=1; s<=seriesCount; s++) {

// Bio-Formats Importer uses an argument that can be built by concatenate a set of strings

run("Bio-Formats Importer", "open=&path autoscale color_mode=Default view=Hyperstack stack_order=XYCZT series_"+s);

original = getTitle();

dotIndex = lastIndexOf(original, ".");

title = substring(original, 0, dotIndex);

print(title);

run("Split Channels");

selectWindow("C1-" + original); //selects channel 1 (in this case, Hoechst staining)

run("Gaussian Blur 3D...", "x=1 y=1 z=1");

C1 = getTitle();

selectWindow("C2-" + original); //selects channel 2 (in this case, MBP staining)

run("Gaussian Blur 3D...", "x=2 y=2 z=2");

C2 = getTitle();

selectWindow("C3-" + original); //selects channel 3 (in this case, alpha-tubulin staining)

run("Gaussian Blur 3D...", "x=2 y=2 z=2");

C3 = getTitle();

run("Merge Channels...", "c1=["+ C1 +"] c2=["+ C2+ "] c3=["+ C3 +"] create");

saveAs("Tiff", out_path + "/" + title + "_Serie_" + s + ".tiff");

close();

}

setBatchMode(false);

**Supplementary pipeline S1** Macro “LIFFtoTIFF.m” from ImageJ version 1.52u 17 March 2020, <https://imagej.nih.gov/ij/notes.html>).

import os, glob, shutil

#os.chdir("C:/Users/alm\_users/Desktop/Cristina\_IMARIS/NocTPR_tiff")

folder = "D:/UsersData/EvaCarvalho_ImgAnalysis/20230830_Exp1U_D9DIFF/TIFFs/OPCsAstrosNoLPS_n1"

for file_path in glob.glob(os.path.join(folder, '*.*')):

new_dir = file_path.rsplit('.', 1)[0]

os.mkdir(os.path.join(folder, new_dir))

shutil.move(file_path, os.path.join(new_dir, os.path.basename(file_path)))

**Supplementary pipeline S2** Script “fileinfolder.py” created with Python 3 <https://www.python.org>.

import pandas as pd

import os

from datetime import date

import seaborn as sns

import matplotlib.pyplot as plt

import json

class IMARISDataProcessor:

""" This class expects a folder generated by IMARIS with the following format

(SELECTED FOLDER)>SampleX>

Series??_cells

Series??_spots

It will concatenate all (series) data correspondent to a single sample type in a single file and

then provide statistics on the distribution on different features extracted:

- intensity of Channel 2

- number of vesicles

- sphericity

- volume

- number of spots

"""

CELLS_SHEET_COLUMN = {

'Vesicles': ['Cell Cytoplasm Number of Ve-3','Cell Cytoplasm Number of Vesicles'],

'Intensity' : ['Cell Intensity Mean Ch=2 Img=1','Cell Intensity Mean'],

'Sphericity' : ['Cell Sphericity', 'Cell Sphericity'],

'Volume' : ['Cell Volume', 'Cell Volume']

}

SPOTS_OUT_COL_NAME = 'Nr. Spots'

def __init__(self, directory, sample_labels = {}):

self.directory = directory

self.ReadConfigFile()

def ReadConfigFile(self):

with open("config_cells.json") as json_data_file:

data=json.load(json_data_file)

self.CELLS_SHEET_COLUMN = data['CellSheetColNames']

self.sample_labels = data['SampleLabels']

self.SPOTS_OUT_COL_NAME = data['SpotOutputName']

self.VESICLES_OVERALL_SHEET = data['VesiclesOverallSheetColNames']

def ExtractSamplesData(self, save_to_excel = True, save_to_pickle = False):

"""ExtractSamplesData returns dataframe containing the samples data

Args:

save_to_excel (bool, optional): Per sample, save an excel sheet per sample containing the respective series data. Defaults to True.

save_to_pickle (bool, optional): Save dataframe containing all the samples data. Defaults to False.

Returns:

pd.DataFrame: Samples data

"""

self.samples_name = self.IdentifySamples()

self.VerifySampleNames()

samples_dataframes = pd.DataFrame()

samples_spots = pd.DataFrame()

\# Iterating over each (sample) folder

for sample in self.samples_name:

print("Processing {}".format(sample))

series = self.IdentifySeries(sample) # and getting all the series

sample_data = pd.DataFrame() # dataframe which will contain all the sample data

sample_spots = pd.DataFrame()

for serie in series:

print("Loading {} ...".format(serie))

\# Handle spots data

nr_spots = self.ExtractSerieSpotsData(sample, serie) # get the cells which we categorized as "spot"

if nr_spots == 0:

print("No Vesicles or No Data|")

continue

serie_spots = pd.DataFrame({'Sample':[sample],self.SPOTS_OUT_COL_NAME:[nr_spots]}, index=[0])

sample_spots = pd.concat([sample_spots,serie_spots])

\# Handle cells data

serie_data = self.ExtractSerieCellsData(sample, serie)

if serie_data.empty:

print("No Vesicles or No Data|")

continue

sample_col_df = pd.DataFrame(

{'Sample':self.CreateColumnForSerie(value=sample,nr_cells=serie_data.shape[0])},

index=serie_data.index.values)

full_serie_data = pd.concat([

sample_col_df,

serie_data],

axis=1)

\# concatenating series data to form the overall sample dataframe

sample_data = pd.concat([sample_data,full_serie_data])

sample_data.reset_index() # remove cell id as index to avoid overriding when concatenating

sample_spots.reset_index()

samples_dataframes = pd.concat((samples_dataframes,sample_data))

samples_spots = pd.concat((samples_spots,sample_spots))

if save_to_excel:

with pd.ExcelWriter(self.directory+sample+'.xlsx') as writer:

sample_data.to_excel(writer,sheet_name="cell_data", header=True)

sample_spots.to_excel(writer,sheet_name="spots_data", header=True)

print("Data saved to {}".format(self.directory))

if samples_dataframes.empty:

return samples_dataframes, samples_spots

samples_dataframes['Sample']=samples_dataframes.apply(self.ReplaceSampleLabels,axis=1)

samples_spots['Sample']=samples_spots.apply(self.ReplaceSampleLabels,axis=1)

if save_to_pickle:

samples_dataframes.to_pickle(self.directory+'cells_data_'+ date.today().strftime("%Y%m%d")+ ".pkl")

samples_spots.to_pickle(self.directory+'spots_data_'+date.today().strftime("%Y%m%d")+ ".pkl")

return samples_dataframes, samples_spots

def IdentifySamples(self):

"""IdentifySamples finds all the samples (each sample has its own folder)

Returns:

[list]: contains sample names

"""

list_samples =[]

entries = os.listdir(self.directory) # all files/folder within provided directory

for entry in entries:

if os.path.isdir(self.directory+entry) and 'Sample' in entry: # if it is a folder and contains "Sample"

list_samples.append(entry) # added to the list of samples

list_samples.sort() # sort them

return list_samples

def VerifySampleNames(self):

if not self.sample_labels:

self.sample_labels= dict(zip(self.samples_name,self.samples_name))

else:

for identified_sample_name in self.samples_name:

if identified_sample_name not in self.sample_labels.keys():

self.sample_labels[identified_sample_name]=identified_sample_name

def IdentifySeries(self, sample_name):

"""IdentifySeries finds within a sample folder, all the series.

def

Returns:

[list]: contains the list of series existing for a given sample

"""

list_series = []

entries = os.listdir(os.path.join(self.directory,sample_name))

for entry in entries:

splitted = os.path.splitext(entry)

if splitted[-1]==".xls" and 'spots' in splitted[0].lower(): # we are expecting per series to exist a "spots" excel

list_series.append(splitted[0].split('_')[0])

list_series.sort()

return list_series

def ExtractSerieSpotsData(self, sample_name, serie_name):

"""ExtractSeriesSpotsData from the provided filename, it extracts the number of spots by checking

, in the "Diameter" sheet, the number of rows of data.

Args:

sample_file ([string]): filename containing the spots data

Returns:

[int]: number of spots in serie

"""

filename = os.path.join(self.directory, sample_name, serie_name)

if (os.path.isfile(filename+'_Spots.xls')):

xls = pd.ExcelFile(filename+'_Spots.xls')

else:

xls = pd.ExcelFile(filename+'_spots.xls')

try:

return pd.read_excel(xls,sheet_name='Diameter',skiprows=1).shape[0]

except:

return 0

def ExtractSerieCellsData(self, sample_name, serie_name):

"""ExtractCellsData extracts serie's cells data from the "Series[XX]_cells.xls" file

Args:

sample_filename ([string]): filename containing the cells data

Returns:

[pd.DataFrame]: dataframe with columns Number of Vesicles, Intensity Mean, Sphericity,

Volume and ID. Only the cells with at least a vesicles were left in the DataFrame

"""

filename = os.path.join(self.directory, sample_name, serie_name)

if (os.path.isfile(filename+'_Cells.xls')):

xls = pd.ExcelFile(filename+'_Cells.xls')

else:

xls = pd.ExcelFile(filename+'_cells.xls')

if not self.CheckIfVesicles(xls):

return pd.DataFrame()

data_dict = {}

for assigned_name, sheet_col_name in self.CELLS_SHEET_COLUMN.items():

data_dict[assigned_name] = pd.read_excel(xls,

sheet_name=sheet_col_name[0],

skiprows=1)[sheet_col_name[1]]

\# data_dict[assigned_name] = df[sheet_col_name[1]].rename(

\#	columns = {sheet_col_name[1]:assigned_name}

\#	)

\# remove nr of vesicles <1

indices = data_dict['Vesicles']>0

sample_data = pd.concat([data[indices.values] for data in data_dict.values()],

axis=1)

for assigned_name, sheet_col_name in self.CELLS_SHEET_COLUMN.items():

sample_data.rename(columns = {sheet_col_name[1]:assigned_name},inplace=True)

sample_data.index.name="Cell ID"

return sample_data

def CheckIfVesicles(self, xls_file):

data = pd.read_excel(xls_file, sheet_name=self.VESICLES_OVERALL_SHEET[0],

skiprows=1)

total = data['Value'].loc[data['Variable']==self.VESICLES_OVERALL_SHEET[1]].values[0]

if total == 0:

return False

return True

def CreateColumnForSerie(self, value, nr_cells):

"""CreateColumnForSerie Generate column nr_cells long containing only value or having only the first cell with value and the remainder filled with zero.

Args:

value ([str, int, float]): value to be added present in the column

nr_cells ([int]): number of cells the column contains

only_first (bool, optional): Whehter the column is filled with the same value or just the first cell bears the value whereas the remainder are set to zero. Defaults to True.

Returns:

[list]: list containing value (in every cell or only the first)

"""

new_col=[value]*nr_cells

return new_col

def ExtractMetricsForSamples(self, cells_df, spots_df, save_to_excel=True):

"""ExtractMetricsForSamples groups dataframe by Sample and extracts statistical information on each of the features.

Args:

df (pd.DataFrame): input dataframe with the full information on the cells

save_to_excel (bool, optional): whether metrics should be exported as an excel file. Defaults to True.

Returns:

[pd.DataFrame]: contains statistics per sample type of the input dataframe

"""

metrics = cells_df.groupby('Sample').describe(percentiles=[])

if save_to_excel:

with pd.ExcelWriter(self.directory+'summary.xlsx',mode = 'w') as writer: # doctest: +SKIP

temp = metrics.unstack(1)[:,'mean'].unstack(0)

sum_vesicles = cells_df.groupby('Sample').sum()['Vesicles']

sum_spots = spots_df.groupby('Sample').sum()[self.SPOTS_OUT_COL_NAME]

sum_vesicles_spots = pd.concat([sum_vesicles, sum_spots],axis=1)

temp['Vesicles'] = sum_vesicles_spots['Vesicles']

temp[self.SPOTS_OUT_COL_NAME] = sum_vesicles_spots[self.SPOTS_OUT_COL_NAME]

temp['%MBP']=temp.apply(self.DetermineMBP, axis=1)

metrics = metrics.drop(columns = 'count', level=1)

metrics = metrics.drop(columns = 'Vesicles')

temp.to_excel(writer, sheet_name= 'summary')

metrics.unstack(1).to_excel(writer, sheet_name="full statistics")

print("Created a summary of the results under {}".format(self.directory+'summary.xlsx'))

return metrics

def DetermineMBP(self,data):

return data['Vesicles']/data[self.SPOTS_OUT_COL_NAME]*100

def ReplaceSampleLabels(self,datarow):

return self.sample_labels[datarow['Sample']]

def GenerateBoxPlot(self,dataframe, feature, x_range = [], swarmplot=True, visualize = False):

if x_range == []:

x_range = [dataframe[feature].min(), dataframe[feature].max()]

f, ax = plt.subplots(figsize=( 20 , len(dataframe['Sample'].unique())*1.5)) # Figure size is set here, you can adjust it

sns.boxplot(x=feature, y="Sample", data=dataframe, palette=sns.light_palette((210, 90, 60), input="husl"))

sns.swarmplot(x=feature, y="Sample", data=dataframe, alpha=0.75, color="0.3")

ax.xaxis.grid(True)

ax.set(ylabel="")

plt.tight_layout()

if visualize:

plt.show()

else:

f.savefig(self.directory+feature+date.today().strftime('%Y%m%d')+".pdf")

return

**Supplementary pipeline S3** Script “restructIMARIS.py” created with Python 3 <https://www.python.org>.
